# Membrane Geometric Confinement Reshapes the Lateral Electric Field Distribution and Intracellular Cargo Transport in Nanopore Electroporation

**DOI:** 10.1101/2025.09.28.679060

**Authors:** Emily McCorkle, Matthew Lee Manion, Xiaoqian Wang, Cheyenne Meeks, Guanren Tao, Sasha Cai Lesher-Pérez, Albert Tianxiang Liu

**Affiliations:** Department of Chemical Engineering, University of Michigan, Ann Arbor, MI 48105; Department of Electrical Engineering, University of Michigan, Ann Arbor, MI 48105; Department of Biomedical Engineering, University of Michigan, Ann Arbor, MI 48105; Biointerfaces Institute, University of Michigan, Ann Arbor, MI, 48109 USA; Department of Material Science and Engineering, University of Michigan, Ann Arbor, MI 48105

**Keywords:** Nanopore electroporation, transfection, device design, dosage control

## Abstract

Nanopore electroporation (NanoEP) is an emerging transfection method that enables efficient and safe intracellular delivery and removal of biomolecular cargo for applications in disease modeling, tissue engineering, and therapeutic biologics manufacturing. Conventional device designs assume uniform vertical cargo flux across nanoporous membranes; however, we demonstrate that the lateral electric field distributions introduce a pronounced edge effect, with enhanced cargo delivery and depletion along the membrane perimeters. We identify and characterize the presence of this edge effect in NanoEP systems, and develop a modified Nernst–Planck model to guide the design of membrane geometries that either promote delivery uniformity or create prescribed spatial gradients within cell monolayers. By varying the internal angles formed by the membrane edges (60°C, 90°C, 120°C), we create predictable intracellular cargo gradients, while concave “serpentine” geometries with high perimeter-to-area ratios amplify delivery efficiency and minimize spatial heterogeneity compared to circular membranes. These findings establish membrane geometry as a tunable design parameter in NanoEP, enabling control over both uniform and patterned intracellular payload delivery or depletion. This geometric design principle offers a scalable strategy for next-generation transfection platforms and synthetic tissue constructs.

## Introduction

Intracellular delivery of bioactive molecules such as nucleic acids, commonly referred to as transfection, is essential for biomedical research and clinical applications, including gene therapy, tissue engineering, and disease modeling^1,2,3,4,5,6^. While numerous transfection techniques exist, they often face trade-offs between efficiency, throughput, and dosage control^7,8^. Broadly, in vitro and ex situ transfection methods fall into two categories: substrate-free and substrate-based. Although substrate-free methods (e.g., viral, chemical, sonoporation, electroporation, etc.) are typically high throughput, they can elicit concerns about cytotoxicity or mutagenicity^7^. Substrate-based transfection techniques have attracted interest for their ability to enhance intracellular delivery while maintaining cell viability through engineered cell–material interfaces^7,9,10,11,12,13,14,15,16,17^. Despite these advantages, substrate-based approaches still struggle to deliver consistent, high-efficiency transfection across entire cell populations.

One promising substrate-based transfection method, nanopore electroporation (NanoEP), employs an insulating membrane substrate (positioned in the horizontal x-y plane) with vertically aligned nanopores to localize the applied electric field along the z-axis (perpendicular to the cells cultured on the membrane substrate)^14,15,18,19^. This localized field forms transient electropores on the cell membrane at the cell–nanopore interface, allowing for precise electrophoretic transport of cargo into and out of cells^14,15^. Unlike bulk electroporation^20^, NanoEP’s substrate-free analog, which exposes suspended cells to high electric potentials (>500 V/mm) and often causes extensive cell damage^21^, NanoEP achieves high delivery efficiency (>60%) and high cell viability (>90%) at much lower voltages (<15 V/mm)^14,15,22,23,24^.

Like most substrate-based transfection techniques, the throughput of NanoEP is thought to scale with the 2-dimensional (2D) area of the nanoporous substrate the target cells interface with––at least in principle. While NanoEP has been primarily studied in the context of its electric field focusing effects along the z-axis, the role of lateral (x-y) electric field distributions in maximizing or controlling delivery remains underexplored. The efficiency and scalability of NanoEP are directly tied to the spatial uniformity of cargo delivery and depletion across a cell population. Because charged cargo transport in NanoEP is dominated by electromigration rather than diffusion^14,24^, non-uniform lateral electric field distribution can significantly impact the spatial uniformity of molecular delivery. Previous studies have shown that electrode geometry can influence field distribution and the associated downstream outcome in brain stimulation^25^, cardiac pacing^25^, and electrodeposition^27^. By increasing the number of edges in an electrode, the current density increases, which leads to a reduction of the electrode impedance^25^. Indeed, the effect of electrode geometry has been explored in the context of NanoEP^28,29^, particularly when the electrode dimensions are significantly smaller than the substrate area.

In reality, as we will report herein, the lateral spatial uniformity of the z-focused electric field in NanoEP cannot be simply assumed, even for devices with large, planar electrodes. We demonstrate that large-area NanoEP systems suffer from a substrate dependent “edge effect”, in which cells near the perimeter of the nanoporous membrane exhibit enhanced delivery and faster cargo depletion than those at the center. This non-uniformity is attributed to a lateral electric field confinement effect created by the geometries of the cell–membrane in tandem with the electrolyte– electrode interfaces. Understanding and controlling these effects are essential for scaling NanoEP-based systems and could enable the creation of not only highly uniform but also intentionally heterogeneous cell populations, which is critical in tissue engineering where spatially controlled gene expression and protein loading is necessary for modeling complex biological systems^30,31^.

In this study, we systematically investigate how device geometry influences lateral electric field distributions in NanoEP. Using both experiments and a theoretical model we developed based on the Nernst–Planck equation, we quantify the observed edge effects and identify design rules to predict kinetic rates for cargo depletion and geometric trends in cargo delivery. Our results demonstrate that by modifying membrane geometry, specifically by introducing angular variations in membrane edges and adjusting perimeter-to-area ratios, we can tune intracellular molecular delivery and depletion gradients within 2D cell populations. These findings suggest that membrane geometry can serve as a tunable design parameter in NanoEP systems to enable both uniform and intentionally patterned intracellular delivery or depletion, offering a scalable strategy for next-generation transfection platforms and synthetic tissue constructs.

## Results

### Edge Effects in NanoEP Devices with Circular Membranes

To examine spatial variations in cargo delivery and depletion during NanoEP, we fabricated devices comprising indium tin oxide (ITO) transparent electrodes, a polydimethylsiloxane (PDMS) chamber containing the cell media or electroporation buffer (which defined the membrane geometry), and a nanoporous track-etched polycarbonate (PCTE) membrane substrate supporting a cell monolayer (**Figure 1a**). In devices with circular membranes, we consistently observed a pronounced *edge effect*, where cargo transport (characterized by its vertical flux) was enhanced at the periphery of the cell monolayer compared to the center, when a perpendicular electric field was applied between the planar ITO electrodes (**Figure 1b-1d**). This effect was evident in both delivery and depletion contexts using HT1080 human epithelial fibrosarcoma cells. Cargo of various sizes, from small molecules (∼700 Da) to larger plasmids (∼5 MDa), exhibited similar trends (**Figure 1b**). Under typical NanoEP delivery conditions, propidium iodide (∼700 Da), a membrane impermeable dye, showed greater cellular uptake at the membrane edge versus the center of the device, as did pLenti3.7-DsRed plasmid (∼5 MDa) stained with YOYO-1 dye. Protein delivery followed the same pattern: cells internalized fluorescently labeled bovine serum albumin (BSA-AF488) with intensities that declined progressively toward the membrane center (**Figure 1b**). BSA-AF488 edge enhanced delivery was also observed in other electroporation buffers such as media and phosphate-buffered saline (PBS) (**Supplemental S1**). Depletion studies, where we tracked calcein removal from HT1080 cells, corroborated these findings. In a 2 mm diameter device, calcein depletion initiated at the periphery and propagated inward, with cells at the edge exhibiting higher depletion rates (*k*) (**Figure 1c**). Because depletion flux measurements decouple cargo concentration gradients from electric field effects, they provide a more direct readout of the field distribution (versus the delivery experiments) and reinforce the notion that electric field non-uniformity, driven by the membrane-edge, governs NanoEP performance.

**Fig. 1:**
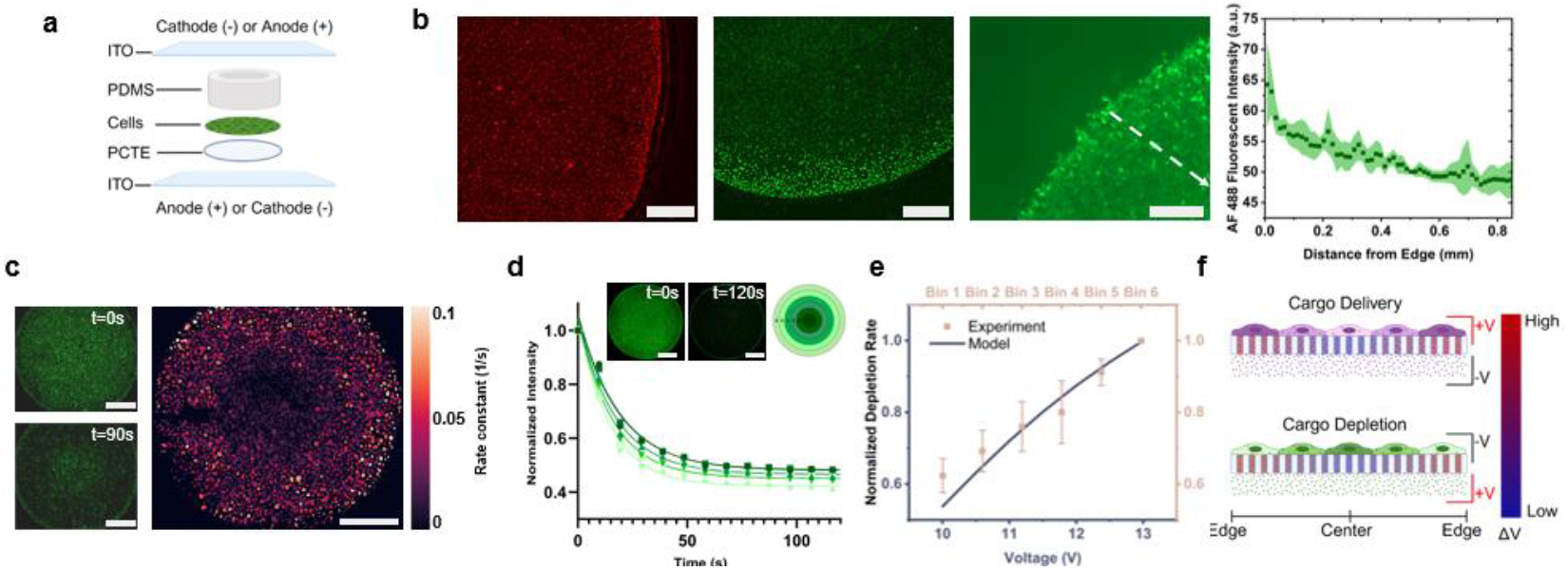
Experimental evidence of edge effects in NanoEP. **a**, Schematic illustration of the NanoEP device used for cargo delivery and depletion experiments. **b**, Left to middle right: representative fluorescent micrographs of HT1080 cells showing NanoEP delivery of: (left) propidium iodide (PI; 1:3 (v/v) PI:electroporation buffer, 20 V, 20 Hz, 1 ms square-wave pulse width, 10 s duration; scale bar, 430 *µ*m), (middle left) YOYO-1–labeled plasmid (1:10 YOYO-1:base pairs, 20 V, 1 Hz, 10 ms square-wave pulse width, 8 s duration; scale bar, 430 *µ*m), and (middle right) Alexa Fluor™ 488–conjugated BSA (BSA-AF488, 1 *µ*g/*µ*L, 25 V, 1 Hz, 10 ms square-wave pulse width, 4 s duration; scale bar, 300 *µ*m). Right: radial decrease in BSA-AF488 fluorescence intensity with distance from membrane edge, *n* = 3 devices. **c**, Left: representative fluorescent micrographs of HT1080 cells showing calcein depletion in a 2 mm diameter circular device before (top) and after (bottom) NanoEP (15 V, 20 Hz, 0.2 ms pulse width, 90 s duration; scale bars, 500 *µ*m). Right: heat map of calcein depletion rate constants across the device (scale bar, 500 *µ*m). **d**, Radial bin analysis of calcein depletion in a 6 mm circular device (15 V, 20 Hz, 1 ms square-wave pulse width, 120 s duration; *n* = 3). Left: normalized intensity decay over time across six concentric bins (Bin 1 = center; Bin 6 = edge). Insets: before/after fluorescent micrographs and schematic of bin regions (scale bars, 1.5 mm). **e**, Normalized rate constants from experimental data across binned regions (points) compared with simulated transport rates across applied voltages (line), capturing the nonlinear dependence of cargo transport on voltage drop. **f**, Schematic illustration of lateral voltage distribution across the PCTE nanoporous membrane during cargo delivery (top) and depletion (bottom), illustrating enhanced cargo flux at the membrane edge.

To investigate the role length scale plays on the observed edge effect, we repeated depletion experiments in a larger 6 mm diameter circular device using 15 V, 20 Hz square-wave pulses with 1 ms pulse widths for 120 s (*n = 3*). To accommodate the lower imaging resolution (as a result of the lower magnification used to fit the entire 6 mm device in one frame), we analyzed fluorescence intensity across six concentric radial zones (referred to as “bins”) to quantify spatial variations. Fluorescence intensity (*I*_*f*_) in each bin was tracked over time until reaching a plateau (*b*) and depletion rate constants (*k*) were extracted by fitting the experimental data to **Equation 1**^28^. A clear gradient emerged from the outermost edge bin (Bin 6) to the innermost center bin (Bin 1), confirming the presence of NanoEP edge effects at larger device scales (**Figure 1d**). Across all circular geometries tested, we consistently observed heightened delivery or removal rates near the edge of the cell monolayer compared to the center, reflecting lateral non-uniformities parallel to the planar ITO electrodes (**Supplemental S2**). Control experiments without applied fields (**Supplemental S3**) verified that these edge effect patterns arose from the applied electric field rather than photobleaching or buffer exposure.

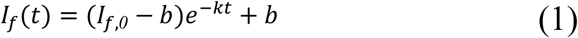

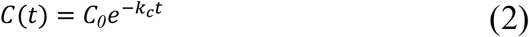

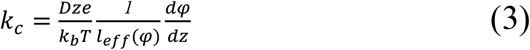

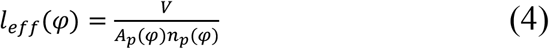

We hypothesized that the observed edge effects stem from lateral non-uniformities in the focused electric field across the nanoporous membrane. To model the observed trends in the cargo transport rate as a function of voltage, we derived **Equation 2**, describing the electrophoretic-driven exponential decay of intracellular calcein concentration (see **Methods**). Molecular diffusion was neglected since the calculated electrophoretic species flux across the nanoporous membrane was ∼1350× greater than that of diffusion (**Supplemental S4)** and dominates cargo transport. Under this theoretical framework (**Equation 3**), the measured rate constant (*k*_*c*_) scales linearly with electric field strength 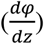, diffusion coefficient (*D*), effective molecular charge (*ze*), electropore area (*A*_*p*_), number of electropores (*n*_*p*_), and inversely with cell volume (*V*) and thermal energy (*k*_*b*_*T*). Due to the presence of a complex, nonlinear dependence of *A*_*p*_ and *n*_*p*_ on the local voltage *φ*^15^, several of these hard-to-estimate parameters were consolidated into a single fitting parameter, *l*_*eff*_, with units of length (**Equation 4**). This term represents an *effective transport length* for charged intracellular cargo exiting through electropores, and its derivation is detailed in **Supplemental S5**.

Using this model, we compared normalized experimental *k* (*k*_*norm*_) values to model *k*_*c,norm*_ predictions across applied voltages (**Figure 1e**), using physically relevant parameters summarized in **Supplemental S4**. Consistent with prior reports^15^, *l*_*eff*_ exhibits a sigmoidal dependence on voltage, and scales linearly with the number of membrane nanopores contacting each cell, itself a function of the commercial membrane’s nanopore diameter. We found that a ∼2–2.5 V voltage differential between the edge and center regions of the cell–membrane monolayer would give rise to a gradient in cargo flux that coincides with the measured normalized rate constants (1 at the edge vs. ∼0.6 at the center, **Figure 1e**), and therefore hypothesized that the enhanced cargo flux at the membrane perimeter stems from a lateral variation in vertical voltage drop across the membrane nanopores (**Figure 1f**). Together, these findings suggest that spatial variation in the electrophoretic driving force, governed by the geometric shape confinement of the nanoporous membrane, could underlie the observed edge effect. This spatial heterogeneity offers a means to create predictable gradients in cargo flux or enhance delivery uniformity by tailoring membrane geometry.

### Electrode-Based Evidence of Radial Electric Field Heterogeneity

To link radial heterogeneity in NanoEP flux with anisotropy in electric field strength, we exploited a well-characterized electrochemical reaction that induces voltage-dependent changes in electrode optical properties, enabling visualization of the lateral field distribution across the device. Specifically, we measured changes in light transmission through the ITO cathodes, which experience reduction currents during NanoEP experiments (**Figure 2a**). At the cathode–electrolyte interface, electrochemical reduction of ITO decomposes its atomic complexes into indium and tin nanoparticles (**Figure 2a**), substantially decreasing optical transparency^32,33^. Because the rate and extent of the ITO reduction, and the corresponding decrease in optical transparency, are known to scale with the local voltage magnitude (i.e., electrochemical overpotential)^32^, changes in transmitted light provide a convenient proxy for mapping electric field distribution across the NanoEP device. To enhance contrast for image analysis, a voltage higher than usual was used. We subjected 2 mm devices with PCTE membranes to 30 V, 20 Hz square-wave pulses (1 ms pulse width) for 40 s (**Figure 2b**). Under these conditions, regions of the ITO near the device perimeter consistently darkened more rapidly than central regions, indicating stronger local electric fields at the edge.

**Fig. 2:**
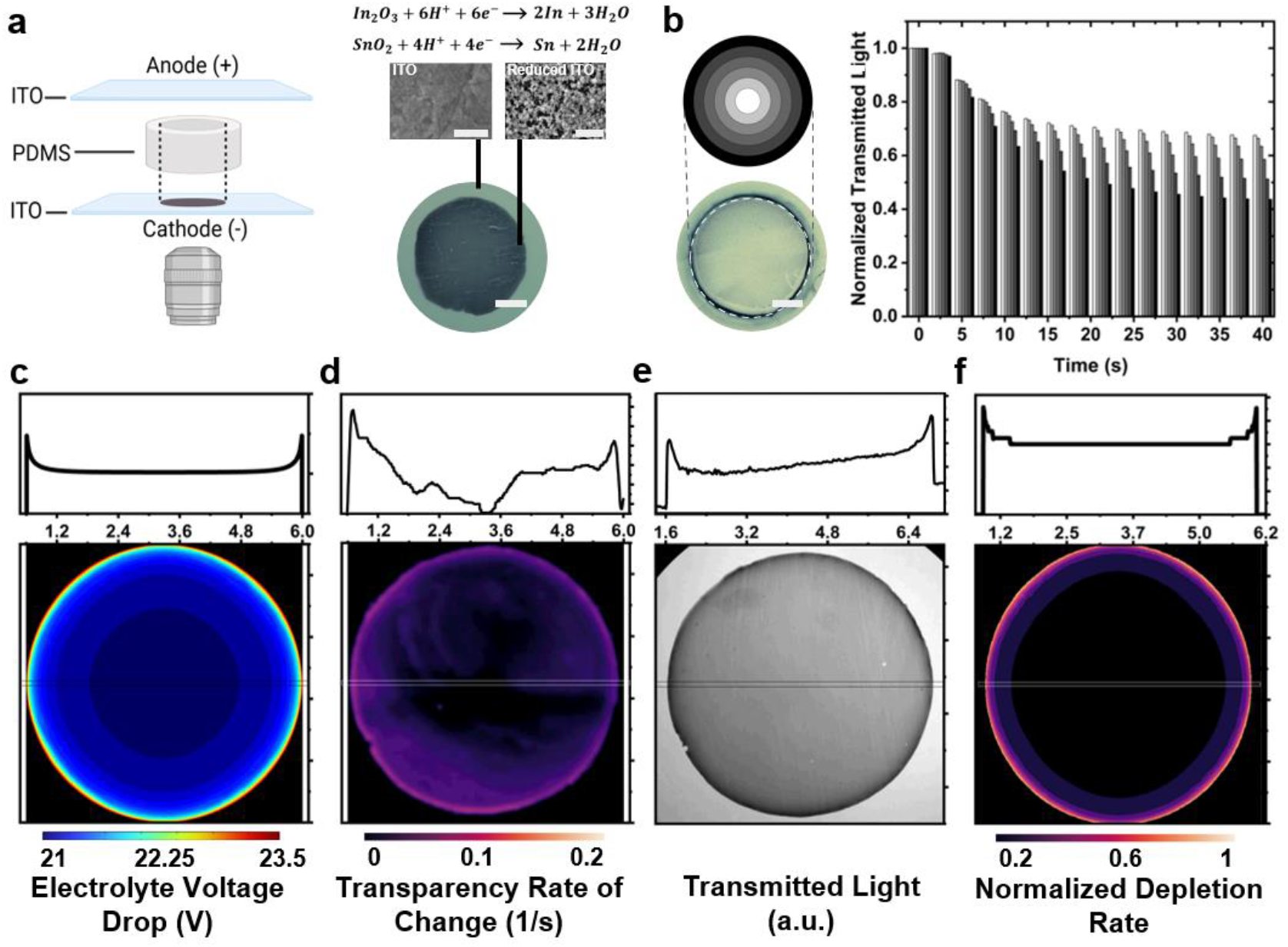
Electrolyte–electrode interfacial reactions reveal lateral voltage heterogeneity in NanoEP devices. **a**, Left: schematic illustration of the device setup for cathodic ITO reduction experiments. Top: pristine ITO film (scale bar, 500 nm). Top right: reduced ITO film post-NanoEP (scale bar, 1 *µ*m). Bottom right: bright field optical micrograph showing darkened cathode following electric stimulation of 2 mm device with PCTE membrane (scale bar, 750 *µ*m). **b**, Transmitted light intensity decreases over time across six concentric radial bins in a 2 mm circular device with PCTE membrane (30 V, 20 Hz, 1 ms square-wave pulse width, 40 s duration; scale bar, 750 *µ*m). **c**, COMSOL-simulated lateral distribution of vertical voltage drop across the electrolyte in a 6 mm circular device with no PCTE membrane under 30 V cathode-to-anode bias. **d**, Heat map of ITO reduction rate constants in a 6 mm circular device with no PCTE membrane under 30 V cathode-to-anode bias (30 V, 20 Hz, 1 ms square-wave pulse width, 40 s duration). **e**, Bright field optical micrograph showing the darkened cathode under 30 V cathode-to-anode bias (same device as 2d). **f**, Heat map of normalized depletion rate constants predicted using the modified Nernst–Planck model. The line profiles above panels c–f represent cross-sections through each 2D heat map, taken along a ∼6 mm slice indicated on the corresponding map.

We note that the edge effect associated with this electrode reaction persisted in the absence of the PCTE membrane. Using a 6 mm device, we simulated the voltage drop across the electrolyte and assessed the rate and extent of reaction experimentally. Both approaches revealed stronger electric fields near the device perimeter. A 3D finite element (COMSOL) simulation of cylindrical NanoEP devices mapped the z-directional voltage drop at each x–y position under a 30 V bias, showing enhanced voltage gradients at device edge compared to the center (**Figure 2c)**. Experimentally, applying 30 V, 20 Hz square-wave pulses (1 ms pulse width, 60 s duration) produced greater optical transparency changes and faster electrode reactions at the edge, consistent with locally elevated field strengths (**Figure 2d, 2e**). Elemental mapping by energy dispersive X-ray spectroscopy (EDX) also confirmed these trends: post-NanoEP EDX maps showed a lower oxygen signal at electrode edges (**Supplemental S6**), consistent with more extensive local ITO reduction. Incorporating the simulated x–y electric field distribution (**Figure 2c**) into our modified Nernst–Planck model (**Equation 3**) predicted higher calcein depletion rate constants at the periphery than the center (**Figure 2f**), in agreement with the cargo flux observed experimentally (**Figure 1c**).

We attribute the emergence of this edge effect to the interplay between the geometry of both the cell–membrane and the electrolyte–electrode interfaces (EEI). Conventional equivalent circuit models of the NanoEP device (**Supplemental S7**) collapse the x–y dimensions into singular lumped elements from electrode to electrode, yielding only one effective voltage drop across the PCTE membrane and cell monolayer^34^. By contrast, finite element simulations of 3D NanoEP devices reveal that when both the EEI extends laterally beyond the cell–membrane interfacial area and when the finite EEI resistance is above a critical threshold (R_ct_ > R_critical_), the vertical voltage drop is amplified at the edges, an effect that disappears when the R_ct_ becomes negligible (R_ct_ < R_critical_) (**Supplemental S7**). Here, R_ct_ represents a lumped interfacial charge transfer resistance (or EEI impedance) to electron transfer and charge storage^35,36^, which depends on factors including voltage, buffer composition, electrode material, and electrode area^35,36^. From these simulations, we infer that charge accumulation in electrode regions not directly contacting the electrolyte drives electronic current along the EEI perimeter and ionic current toward the edge of the nanoporous membrane, which represents the paths of least resistance. These combined currents elevate local voltages across the cell–membrane interface, thereby enhancing cargo flux during NanoEP, assuming a laterally-uniform vertical transport resistance. Therefore, accounting for lateral non-uniformities is necessary in NanoEP models. Additional considerations of spatial resistance variations are provided in **Supplemental S8**. Taken together, these modeling, optical, and chemical data provide compelling evidence that membrane geometry induces lateral (radial) gradients in the electric field during NanoEP, in situations where the electrode surface area extends beyond the membrane edges. These field gradients, in turn, explain the observed non-uniform NanoEP cargo flux and support the idea that the observed edge effect is controlled by the geometry of both the cell–membrane and electrolyte–electrode interfaces.

### Creating Distinct Gradient Profiles with Angled Membrane Geometry

Building on our theoretical framework, we explored how deviations from circular geometries, specifically the introduction of membrane corners (i.e., locations where two edges meet), affect spatial distribution of cargo flux during NanoEP. We hypothesized that electric field amplification at corners would generate distinct transport profiles compared to smooth circular boundaries. To test this, we designed membrane geometries with varying internal angles between two adjacent edges (60°C, 90°C, 120°C).

Calcein depletion experiments were performed under identical square-wave pulsing (15 V, 20 Hz, 1 ms pulse width, 120 s duration) to evaluate the spatial dependence in field-driven molecular transport, with device height and membrane surface area matched to the 6 mm circular control (**Figure 1d**). This ensures comparable impedances between the two electrodes, which is the summed contribution of EEI charge transfer resistance, solution resistance, and membrane ion transport resistance across the nanopores (**Supplemental S7**). Depletion was imaged in real time through the anode to avoid optical interference from the cathode reduction (**Figure 3a**), and depletion rate constants from the cell monolayers were extracted from the exponential fluorescence decay kinetics for each geometry (**Figure 3b**). Representative fluorescent micrographs before and after NanoEP for each device geometry can be found in **Supplemental S9**. COMSOL simulations of the x–y electric field distribution were performed for each geometry, and the resulting voltage profiles were inputted into our modified Nernst–Planck model to calculate depletion rate patterns (**Figure 3c**). Our simulations predicted that the sharpest corner (60°C) would exhibit the highest local depletion rates, with the highest gradient extending inward from the corner. Experimental calcein depletion data validated these predictions, as shown in representative rate constant heatmaps for each geometry (**Figure 3d**).

**Fig. 3:**
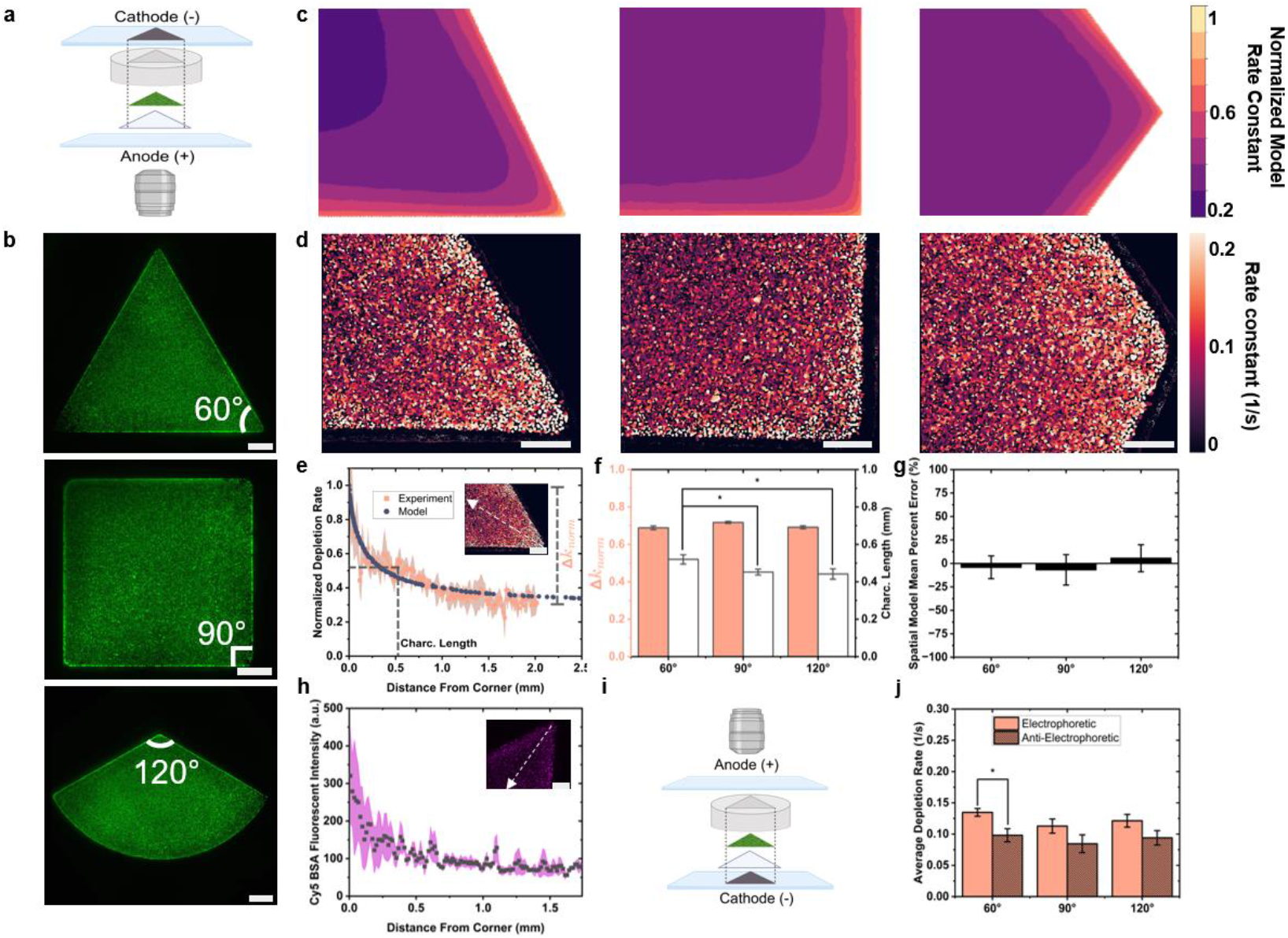
Validation of the NanoEP transport model across membrane geometries. **a**, Schematic illustration of NanoEP device setup and imaging orientation for electrophoretically driven cargo transport. **b**, Full-device fluorescence micrographs of Calcein AM–stained HT1080 cells in triangular (60°C), square (90°C), and obtuse (120°C) geometries (scale bars, 1 mm). **c**, Normalized model rate constant heat maps for 60°C, 90°C, and 120°C NanoEP devices. **d**, Representative experimental rate constant heat maps of calcein depletion for 60°C, 90°C, 120°C NanoEP devices (scale bars, 430 *µ*m). **e**, Comparison of simulated and experimental normalized rate constants as a function of distance from corner for 60°C NanoEP devices (*n* = 3; scale bar, 430 *µ*m). **f**, Experimental Δ*k*_*norm*_ (left) and characteristic length (right) for each device geometry tested (*n* =3). **g**, Spatial mean percent error between model and experiment for each geometry, where each point *i* is a distance coordinate from the device corner. **h**, Fluorescence intensity profiles of BSA-AF647 (2.5 mg/mL) delivery as a function of distance from a 60°C device corner under 20 V, 20 Hz, 1 ms square-wave pulse width, 10 s duration (*n* = 3; scale bar, 430 *µ*m). **i**, Schematic illustration of NanoEP device setup and imaging orientation for anti-electrophoretically driven cargo transport. **j**, Average calcein depletion rate constants under electrophoretic vs. anti-electrophoretic device configurations (*n* = 3 per device geometry).

To quantify lateral gradients in cargo transport, we compared simulated and experimental (*n* = 3) calcein depletion rate constants as a function of distance from membrane corners. The Nernst– Planck model predictions showed strong quantitative agreement with experimental results when mapped to the simulated voltage profile for each geometry (**Figure 3e**, refer to **Supplemental S10** for 90°C and 120°C). Our model was calibrated using independent datasets collected from corner and center regions, corresponding to the line plot endpoints (**Figure 3e**), with training and evaluation data acquired on different days. Calcein depletion rates were measured at each lateral spatial location and fit to an exponential decay function (**Equation 1**). With rate constants expressed as a function of distance from the corner, we defined a characteristic length (Char. Length, **Figure 3e**): the distance at which the change in normalized rate constants (Δ*k*_*norm*_ = *k*_*corner*_ – *k*_*center*_) decayed to 1/e of its initial value ((Δ*k*_*norm*_)/*e* + *k*_*center*_), where *k*_*corner*_ is the depletion rate constant evaluated at the device corner, *k*_*center*_ is the asymptote rate constant, and *e* is Euler’s number). Both simulations and experiments revealed that sharper angles produced longer characteristic lengths (**Figure 3f**). This trend is intuitive: acute angles (e.g., 60°C) keep cells in close proximity to one or more device edges, thereby sustaining an enhanced “edge effect” over a greater distance from the corner. In contrast, wider angles (e.g., 120°C) more closely resemble circular geometries, resulting in a shorter range over which the enhanced electric field persists, hence a shorter characteristic length. We also compared the Δ*k*_*norm*_ across geometries. While our model predicted larger differences in sharper angles, experimental data showed no significant variation in Δ*k*_*norm*_ between angles. Importantly, simulated rate constants at all lateral positions fell within the 95% confidence interval of experimental values, indicating strong overall agreement (≤ 12% average error, **Figure 3g, Supplemental S10)**. Positive mean percent error 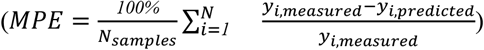 indicated model underprediction for large-angle geometries, whereas negative MPE values for acute angles (≤ 90°C) suggested overestimation of the lateral electric field enhancement. Overall, these results demonstrate that our model can capture cytoplasmic calcein transport driven by NanoEP across a range of confined membrane geometries, while also highlighting subtle deviations in how lateral field enhancement scales with the corner angle.

To extend these findings to delivery applications, we conducted analogous geometric studies using Alexa Fluor™ 647–conjugated bovine serum albumin (BSA-AF647) as a model protein cargo. We ensured uniform cargo distribution beneath the nanoporous membrane during NanoEP delivery, by incorporating 200 *µ*m-tall PDMS pillars to elevate the membrane from the bottom electrode (**Supplemental S11**). Similar to the depletion experiments, fluorescent intensity was enhanced in cells near the corner of the triangular (60°C) devices and decayed nonlinearly toward the center (**Figure 3h, Supplemental S12**). The shape of the normalized BSA-AF647 fluorescence closely matched that of normalized calcein depletion in the same geometry (**Supplemental S13**), underscoring the generality of geometry-induced transport gradients across both delivery and depletion modalities. However, their characteristic lengths differed (0.275 mm for BSA delivery vs. 0.520 mm for calcein depletion), as did Δ*k*_*norm*_ (0.78 for BSA vs. 0.69 for calcein). The shorter characteristic length and larger Δ*k*_*norm*_ for BSA likely reflect second-order effects of its larger molecular size (67 kDa for BSA vs. 0.6 kDa for calcein), though direct comparison is complicated by differences in how cargo transport was quantified (rate vs. amount).

We next examined how electrode polarity influenced calcein depletion across geometries. As before, imaging was always performed through the anode to avoid issues associated with cathodic reaction (**Figure 3a** and **3i**). As expected for a (negatively) charged molecule, calcein exhibited faster average depletion rates in the electrophoretic (anode-downward, **Figure 3a**) configuration than when polarity was reversed (anti-electrophoretic, **Figure 3i**). Importantly, the edge effect persisted in angled devices for both configurations, with corners consistently showing faster depletion than centers (**Supplemental S10**). We attribute this to elevated transmembrane potentials (across the cell–membrane interface) at the edges, which promote more and larger electropores on cells within the applied voltage range^5^. Both the transmembrane potential (governing electropore formation) and the potential across the PCTE membrane (driving cargo flux) contribute to the observed edge effect. Together, these findings validate that the heightened electric field at the edge of the EEI interface accurately translates into predictable spatial flux patterns for both cargo depletion and delivery.

### Increasing Device Perimeter-to-Area Ratio to Enhance NanoEP Delivery Efficiency and Reduce Lateral Variation

Beyond creating device geometries for heterogeneous cell cargo manipulations, we sought to design a membrane shape that minimizes lateral variations in NanoEP-mediated delivery while enhancing overall flux, guided by our theoretical model. Conventional NanoEP devices typically employ circular wells^14,15,24,28,29,34,37,38,39^, which are convex and characterized by low perimeter-to-area ratios. In such designs, most cells reside far from the membrane edge and thus do not benefit from edge-enhanced field-driven transport, a limitation that becomes more pronounced when scaling up NanoEP systems for larger cell populations.

To overcome this, we developed a concave, “serpentine” membrane geometry (**Figure 4a**) with a perimeter-to-area ratio of 2.03, compared to 0.67 for a 6 mm circular device of equal area. By positioning more cells in proximity to the membrane edge, we hypothesized that the serpentine design would amplify the edge effect, thereby boosting intracellular depletion and delivery efficiency while reducing lateral heterogeneity across the cell monolayer.

**Fig. 4:**
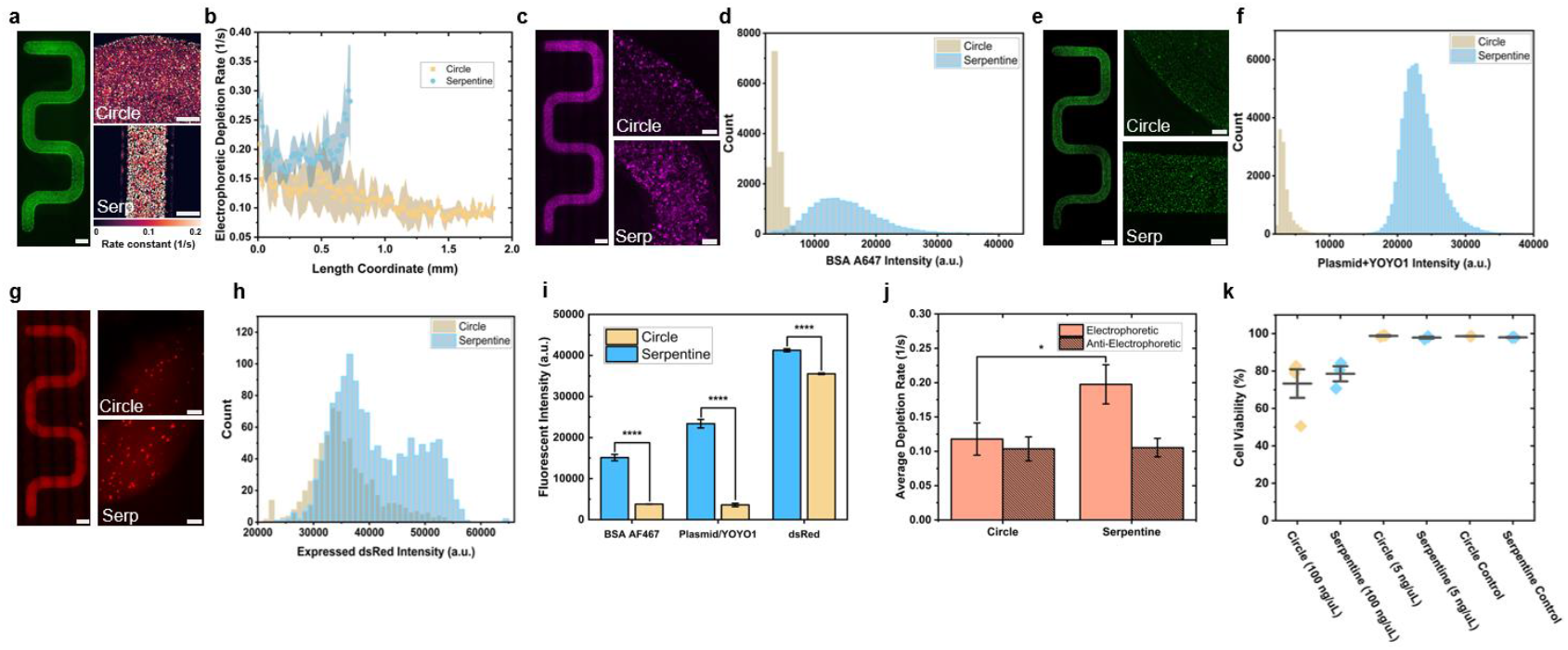
Increasing device perimeter-to-area ratio enhances NanoEP-mediated cargo depletion and delivery. **a**, Left: full-device fluorescent micrograph of Calcein AM–stained HT1080 cells in serpentine device geometry (scale bar, 1 mm). Right: rate constant heat maps of calcein depletion in circular (top) vs. serpentine (bottom) geometries at 15 V, 20 Hz, 1 ms square-wave pulses, 120 s duration (scale bars, 430 *µ*m). **b**, Comparison of depletion rate constant as a function of distance from the edge for serpentine vs. circular devices (*n* = 3 per geometry). **c**, Left: full-device fluorescent micrograph of a serpentine device following BSA-AF647 delivery (2.5 mg/mL; scale bar, 1 mm). Right: representative fluorescent micrographs for BSA-AF647 delivery (2.5 mg/mL) in circular (top) and serpentine (bottom) devices at 20 V, 20 Hz, 1 ms square-wave pulse width, 10 s duration (*n* = 3 per geometry; scale bars, 200 *µ*m). **d**, Histogram of single-cell BSA-AF647 fluorescence intensities in circular vs. serpentine geometries. **e**, Left: full-device fluorescent micrograph of a serpentine device following plasmid (pLenti3.7-DsRed, ∼5 MDa, 100 ng/*µ*L, YOYO-1 labeled) delivery (scale bar, 1 mm). Right: representative fluorescent micrographs in circular (top) vs. serpentine (bottom) devices at 25 V, 1Hz, 10 ms square-wave pulse width, 4 s duration (scale bars, 200 *µ*m). **f**, Histogram of single-cell plasmid+YOYO-1 fluorescence intensities per cell in circular vs. serpentine geometries. **g**, Left: full-device fluorescent micrograph of a serpentine device showing DsRed protein expression 48 h after plasmid delivery (pLenti3.7-DsRed, 5 ng/*µ*L; scale bar, 1 mm). Right: representative fluorescent micrographs for DsRed expression in circular (top) vs. serpentine (bottom) devices at 25 V, 1 Hz, 10 ms square-wave pulse width, 4 s duration (scale bars, 200 *µ*m). **h**, Histogram of single-cell DsRed fluorescence intensities in circular vs. serpentine geometries. **i**, Summary of fold-changes in fluorescent intensity per cell for BSA-AF647, plasmid+YOYO-1, and DsRed expression in circular vs. serpentine geometries. **j**, Average depletion rate constant for electrophoretically vs. anti-electrophoretically driven cargo transport in circular vs. serpentine geometries. **k**, Cell viability 24 h (100 ng/*µ*L samples) and 48 h (5 ng/*µ*L and control samples) post-transfection for plasmid+YOYO-1 delivery experiments.

We compared experimental calcein depletion rates in serpentine versus 6 mm circular geometries (**Figure 4a, 4b**; *n* = 3, **Supplemental S14**). The serpentine rate constant profile followed a symmetric U-shaped curve, with a maximum normalized rate difference (Δ*k*_*norm*_) of ∼0.47, representing the most uniform depletion observed across all geometries tested. This improvement is attributed to the serpentine’s parallel dual-edge configuration, which places more cells near regions of elevated electric field by minimizing the distance of a cell to an edge. By contrast, circular geometries showed a monotonic decrease in rate constants with distance from the edge. Moreover, serpentine devices exhibited higher overall depletion rates, consistent with stronger local electric fields.

We next assessed BSA-AF647 protein delivery across circular and serpentine devices (**Figure 4c**). Population-level fluorescence analysis revealed higher overall intensity in the serpentine geometry compared to the circle (**Figure 4d, Supplemental S15**). Variability in the serpentine device was attributed to local differences in cell adhesion and confluency on the PCTE nanoporous membrane, whereas circular devices displayed a systemic radial gradient of decreasing fluorescence (indicative of amount of cargo delivered) with distance from the edge (**Supplemental S16, S17**). Histogram analysis of single-cell fluorescence intensities (**Figure 4d, Supplemental S15**) confirmed a > 3-fold higher mean fluorescence intensity in serpentine devices compared to circles (15,128 vs. 3,787 a.u.). These results demonstrate that employing concave geometries with high perimeter-to-area ratios effectively amplifies edge-enhanced NanoEP flux, thereby improving protein delivery efficiency while minimizing spatial heterogeneity at the device level.

To evaluate edge-enhanced delivery of larger molecular cargos, we delivered a plasmid encoding DsRed fluorescent protein (pLenti3.7-DsRed plasmid, ∼5 MDa, 100 ng/*µ*L) labelled with YOYO-1 (**Figure 4e, Supplemental S15, S16, S18**). A PCR purification kit was used to remove unbound YOYO-1, ensuring that virtually all measured fluorescence came from the plasmid-bound dye (**Supplemental S19**). Under these conditions, the serpentine geometry showed markedly higher single-cell fluorescent intensity and a greater number of transfected cells to the circular geometry (**Figure 4f, Supplemental S15**). The spatial distribution again followed a U-shaped delivery profile (**Supplemental S16, S18**), with random local fluctuations likely driven by differences in cell coverage, consistent with trends observed for the BSA-AF647 protein cargo. The high plasmid dose used in these experiments facilitated visualization but precluded assessment of protein expression due to plasmid-induced toxicity^14,40^. For functional delivery, we repeated the pLenti3.7-DsRed plasmid delivery experiments at a lower plasmid concentration (5 ng/*µ*L) and assessed DsRed expression 48 hours post-transfection (**Figure 4g**). Single-cell fluorescent analysis confirmed both a greater proportion of DsRed expressing cells and higher mean expression levels in the serpentine devices (**Figure 4h, Supplemental S15**). Cell viability remained > 95% after 48 hours at this lower dose and was comparable to untreated controls (**Figure 4k**). At higher concentrations (100 ng/*µ*L), cells began to come off the nanoporous membrane 24 hours post transfection, so viability was taken at 24 hours rather than 48 hours post transfection to minimize the extent of cell loss.

Aggregated data across different cargos show a consistent trend of higher delivery or depletion in the serpentine compared to the circular devices (**Figure 4i, j**): ∼2-fold higher calcein depletion rates, ∼4-fold higher BSA-AF647 fluorescent protein delivery, ∼7-fold higher plasmid delivery, and ∼1.2-fold higher DsRed expression. These findings further support the hypothesis that devices employing concave membrane shapes with higher perimeter-to-area ratios substantially improve NanoEP-mediated macromolecular transport efficiency and spatial uniformity.

### Understanding the Role of Membrane Geometry in Electric Field Distribution and Cargo Transport Control

Through combined experimental and simulation-based studies, we demonstrated that lateral nanoporous membrane geometry directly influences vertical NanoEP flux (along the z-axis) by reshaping the in-plane (x-y) electric field distribution across the membrane nanopores. By controlling this lateral field distribution, we can in turn modulate the spatial uniformity of intracellular cargo delivery and depletion across a cell monolayer. Using a modified Nernst–Planck molecular flux model, we accurately predicted cargo transport rates (e.g., depletion kinetics) from device-specific lateral voltage profiles by introducing an effective transport length (*l*_*eff*_), defined as the ratio of cell volume to total electropore area, which is a function of local electric fields. Conceptually, *l*_*eff*_ represents an ensemble-averaged path length for cargo molecules leaving the cell: smaller *l*_*eff*_ values correspond to faster depletion rates. This modeling framework enables rational design of membrane geometries to achieve targeted intracellular delivery or depletion profiles.

We found that *l*_*eff*_ must be tuned for different sub-regions of the cell–membrane interface to accurately model cargo transport. Subdividing the nanoporous membrane into lateral “corner” and “center” zones substantially improved model fidelity. As expected from the sigmoidal dependence of *l*_*eff*_ on local voltage (*φ*), which reflects the voltage-driven increase in the electropore number (*n*_*p*_) and area (*A*_*p*_) in cells during NanoEP, *l*_*eff*_ was lowest at device corners. Specifically, we calculated *l*_*eff*_ values of 2.33 mm (center) and 0.92 mm (corner), confirming substantially enhanced field-driven cargo transport near corners of the nanoporous membrane. While these values exceed the dimension of a single cell, they are empirical measures that likely include transport retardation effects from thermal fluctuation (molecular random walk) and electropore cycling (opening/closing). Cells were pulsed with a 2–4% duty cycle at 20 Hz (50 ms period), and prior studies suggest large electropores close within ∼100 *µ*s^15^. Thus, because 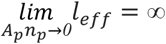, and the field is “off” more than “on”, measured *l*_*eff*_ values are higher than physically intuitive. Using the expression 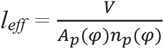, with an estimated HT1080 cell volume V = 2 × 10^−15^ m^3^, electropore radius *r*_*p*_ = 15 nm (*A*_*p*_ = 706 nm^2^), we estimated 1212–3065 electropores per cell, with the highest density near corners. Given ∼ 3500 nanopores underneath each cell (nanopore diameter = 200 nm), these estimates suggest 35–88% of the nanopores within the PCTE membrane actively contribute to intracellular cargo transport in a confluent cell monolayer, assuming one cell electropore per membrane nanopore. In practice, a single nanopore may host multiple electropores of varying radii^15^.

Optimizing plasmid delivery via NanoEP has been a major focus of research over the past decade^8,14,15,24,28,29,39,41^. Compared to protein delivery, plasmid transfection presents unique challenges because successful gene expression requires uptake and an optimal intracellular copy number, enough to enable detectable expression without inducing toxic overexpression. Here, we show that membrane geometry, an often-overlooked aspect of NanoEP device design, can profoundly influence both delivery and expression outcomes, introducing a hidden layer of variability unless carefully controlled. Designing patterned PCTE substrates that enhance uniformity of cargo delivery while promoting scalable transfection is therefore of critical importance. While our serpentine geometry highlighted the impact of edge-enhancing device structures, its complex shape is not readily scalable for manufacturing. Interestingly, several prior reports have inadvertently validated our edge effect hypothesis using single-cell NanoEP devices with patterned silicon membranes^42^ or photoresist-patterned PCTE substrates^43^. In both cases, as in our serpentine design, cells are positioned adjacent to an “edge”, which acts as a high-resistance barrier to the electric field and mass transport. This boundary concentrates the electric field and flux in nearby regions of lower resistance, amplifying localized cargo transport. These insights point toward a generalizable design principle: patterning nanoporous membranes to strategically increase edge-adjacent areas. Moving forward, we aim to investigate the optimal void fraction and feature size of patterned PCTE substrates to facilitate uniform NanoEP transfection across large contiguous areas (e.g., T25 flask scale); a feat which up till now has not been accomplished in the literature.

Although the electric field is the dominant factor shaping cargo delivery profiles, other factors not captured by our COMSOL or Nernst–Planck model may contribute. For instance, electroosmotic flow can impose pressures up to ∼1 kPa on the cell monolayer during electroporation^37^, which may oppose the migration of negatively charged cargo (e.g., calcein) toward the anode (+), particularly in the device center. Additional system-level variability arises from the stochastic distribution of nanopores in PCTE membranes and from non-uniform cell coverage or membrane contact, all of which can introduce noise and heterogeneity in cargo flux. Despite these challenges, we successfully demonstrated that lateral electric field distributions, and thus intracellular NanoEP delivery or depletion outcomes, can be predicted and customized simply by altering membrane geometry. Breaking lateral symmetry enables device designs that promote either more uniform or intentionally patterned delivery. This geometric control offers a viable path toward scalable, substrate-based transfection platforms for clinically relevant applications, including complex 2D tissue models. While further work is required to identify scalability limits, increasing the perimeter-to-area ratio remains a promising strategy for expanding effective substrate surface area and enhancing delivery performance.

## Conclusions

In this study, we demonstrated that NanoEP membrane geometry plays a critical role in shaping the in-plane (x-y) electric field distribution and, consequently, drives anisotropic molecular flux during cargo delivery or removal. By systematically varying nanoporous membrane geometries, we showed that the spatial heterogeneity of cargo transport can be predictably controlled through lateral electric field manipulation. We identified and characterized a geometry-dependent edge effect, wherein enhanced cargo transport occurs near the perimeter of the nanoporous membrane due to lateral gradients of the vertically amplified electric field. By coupling experimental results with finite element voltage simulations and a modified Nernst–Planck flux model, we established strong agreement between predicted and observed lateral heterogeneity in cargo flux, validating the underlying electrokinetic mechanism. Building on this insight, we introduced device geometries with various internal angles to guide spatial gradients in both cargo delivery and depletion. We further demonstrated that concave geometries with high perimeter-to-area ratios significantly reduce the average distance between target cells and membrane edges, leading to higher efficiency and more uniform delivery of protein and plasmid cargos across the cell monolayer. Overall, this work establishes the previously overlooked lateral membrane geometry as an important design parameter for controlling localized electric fields and the resulting intercellular cargo distribution in substrate-based delivery systems.

## Acknowledgements

Schematic figures were generated in BioRender. HT1080 cells were provided to us from the Thurber lab at the University of Michigan. SEM images were taken at the Michigan Center for Materials Characterization (MC^2^). We would like to thank the University of Michigan College of Engineering START grant (Grant Number: U081613). We would also like to acknowledge support from the American Heart Association under Award No. 25IPA1455592 (https://doi.org/10.58275/AHA.25IPA1455592.pc.gr.235709), E.M. through the NSF GRFP under Grant No. DGE 2241144. In addition, A.T.L. would like to acknowledge supports from the National Science Foundation (Grant Number: 2243104, Center for Complex Particle Systems, COMPASS), American Chemical Society Petroleum Research Fund (Grant Number: 66979-DNI10), the Michigan Materials Research Institute (MMRI), and the COMPASS-Biointerfaces Institute Challenge Award. We would like to thank Bobby Kent from the Baker Lab at the University of Michigan for assistance with the MATLAB code for creating a mask around the cells for image analysis.

## Methods

### Device Fabrication

Molds of different geometries were designed with Fusion 360 and 3D printed with the Asiga Pro 4k45 DLP printer. DentaMODEL (Asiga) resin was used for the printing material. Post-printing, the 3D printed molds were cured with UV light for 2 minutes at 36 W (Asiga Flash), followed by a 30-minute isopropyl alcohol wash, and then baked overnight at 60 °CC. The area of the device geometries was kept constant to 28.3 mm^2^ as well as a height of 2 mm unless specified otherwise. Polydimethylsiloxane (PDMS Sylgard™ 184 DOW) at a 10:1 ratio (base to curing agent) was cast into the molds and cured overnight at 60 °CC. To attach the polycarbonate track etched membranes (PCTE) of 200 nm (calcein depletion, propidium iodide and BSA delivery) or 800 nm (plasmid delivery) pore diameters (Cytiva Whatman) to the PDMS devices, uncured PDMS was spun coat on a glass slide to create a 20 *µ*m PDMS layer. Devices were then tapped onto the glass slide to coat the device surface with uncured PDMS, and the PCTE membranes were then placed on the device and cured overnight at 60 °CC^24,44^. Indium tin oxide (ITO) glass slides (Nanocs, 5 Ω /sq) were used as both the positive and negative electrodes.

### Cell Culture and Seeding in Devices

HT1080 cells, a fibrosarcoma cell line, were cultured in ATCC Eagle’s Minimum Essential Medium (EMEM) with 10% fetal bovine serum (Gibco) and 1% penicillin streptomycin (Gibco). On Day 0, the devices were treated with oxygen plasma for 1 minute (Plasma Etch, Inc.) for sterilization and increasing wettability. The devices were then coated with 27.5 *µ*g/mL of fibronectin, from human plasma (Sigma-Aldrich), in phosphate buffered saline (Gibco) and incubated for 1 hour. Following fibronectin incubation, the devices were washed with media and HT1080 cells were seeded in the devices at a cell density of 125,000 cells/cm^2^. Cells were electroporated the following day.

### Electroporation: Cargo Delivery and Depletion

All cells were electroporated one day after cell seeding (Day 1) with Gene Pulser Electroporation Buffer (Bio-Rad). Cells were exposed to the buffer for roughly 10 minutes. A VSP-300 Potentiostat (BioLogic) was used for applying the electrical pulses. The electroporation buffer or cargo solution was pipetted on the bottom electrode at a volume of 10-100 *µ*L. The device was then placed on top of the droplet and overfilled with electroporation buffer to prevent any air gaps when placing the top electrode above the device^15,24^. Copper tape was used to connect the alligator clips to the ITO slides.

For delivery of Bovine Serum Albumin (BSA) Alexa Fluor™ 488 and 647 conjugates (Invitrogen™), the cells were pre-stained with Hoechst 33342 (Invitrogen) at 15 *µ*g/mL, and fluorescent BSA cargo solution was placed at a concentration of 2.5 mg/mL in the electroporation buffer. The cells were electroporated at an applied voltage of 15-30 V at 20 Hz and 1 ms pulse square wave pulse widths for 5-20 s. The top electrode was the anode (+) and the bottom electrode was the cathode (-). BSA delivery images were taken approximately 15 min after electroporation on the ECHO Revolve microscope or the Cytation 5 (Biotek Agilent).

For depletion of calcein, cells were pre-stained with calcein AM (Invitrogen) at 3 *µ*M for 30 minutes, approximately 1 hour before electroporation. The cells were electroporated at an applied voltage of 15 V at 20 Hz and 0.2-1 ms pulse widths for 60-120 s. For electrophoretically driven calcein removal, the top electrode was the cathode (-) and the bottom electrode was the anode (+), imaging through the nanoporous membrane to avoid imaging the cathode reduction. For anti-electrophoretically driven calcein removal, the top electrode was the anode (+) and the bottom electrode was the cathode (-), imaging above the nanoporous membrane to avoid imaging the cathode reduction. Calcein depletion experiments were recorded in real time on the ECHO Revolve fluorescent microscope.

For delivery of plasmid, pLenti3.7-DsRed plasmid (5 MDa) was stained with YOYO-1 dye (Biotium) at a ratio of 10 base pairs: 1 YOYO-1 molecule. The solution was left to incubate at 37°CC for 2 hours. A PCR purification kit (GeneJET, Thermo Scientific) was used to separate unbound YOYO-1 molecules from the solution of bound YOYO-1 molecules to the plasmid. Solutions were diluted with an electroporation buffer at concentrations of 5 to 100 ng/*µ*L. The cells were electroporated at an applied voltage of 25 V, 1 Hz, 10 ms square-wave pulses for 4 s duration. Cell viability was taken 24–48 hours after electroporation with Cy5 NucSpot® Nuclear Stain (Biotium) and Hoechst 33342 (Invitrogen).

### ITO Reaction Experiment

For imaging the reaction of the cathode, 2 mm and 6 mm diameter PDMS well devices (2 mm tall) were used with or without a membrane with an applied voltage of 30 V, 20 Hz, 1 ms square-wave pulses for 40–60 s duration. No cells were used, and Gene Pulser Electroporation Buffer was used as the buffer. Images were analyzed using MATLAB or Python to measure the reaction over time.

### Potentiostat Data Collection

Biologic’s EC Lab software was used to control the electrical parameters of the NanoEP experiments. Prior to running the electroporation experiment, potentio-electrical impedance spectroscopy (PEIS) was conducted over frequencies ranging from 2 MHz to 500 mHz, with a sine amplitude of 10 mV and base potential of 0 V DC. No reference electrodes were used in this study. PEIS was used to elucidate the solution/device resistance near 100-1 kHz, the *x*-coordinate where the imaginary (y-coordinate) Nyquist impedance is ∼0 Ω. PEIS was also conducted after the NanoEP experiment to see if the device resistance changed. NanoEP was conducted using the software’s differential pulse amperometry (DPA) technique. Average currents reported are only when the pulse is applied, not an average of the on and off state.

### COMSOL Simulation & Circuit Modeling

The COMSOL geometry was created and evaluated using the Primary Current Distribution package. All simulations were run at 37 °C. The electrolyte was modeled as water, with a fixed conductivity of 0.2 S/m. Electrodes for the 3-D geometry were ITO and 100 nm thick, with a conductivity of 3 × 10^6^ S/m. The simulation incorporated a “surface resistance” node at the EEI, with an experimentally measured value of 0.0015 Ωm^2^. This value was similar for all tested device geometries. The surface resistance was experimentally determined by calculating the average voltage drop over the electrolyte–electrode interface (EEI; averaged among several devices of a certain shape), dividing by the current through the system, and multiplying it by the surface area. In all simulations, both electrodes were matched to have the same surface resistance and capacitance. The top electrode is separated from the bottom by ∼2 mm. Equivalent circuit modeling was performed using the LTspice software. Built-in modules for circuit elements were used, and a square pulse element was used to apply the voltage. No product specifications were added to the circuit elements.

### SEM Imaging

SEM imaging was performed using the Thermo Fisher Nova 200 Nanolab at the Michigan Center for Materials Characterization (MC^2^). The ITO coated glass was mounted using conductive carbon tape and grounded using colloidal graphite glue (Electron Microscopy Sciences). Images of ITO surfaces were acquired using secondary electrons at 5 kV and 1.6 nA of beam current.

### Nernst-Planck Simulation

To leverage simulated voltage profiles to easily predict cargo depletion rates, a simplified Nernst– Planck (NP) equation was implemented in Python. The movement of charged molecules in solution is described by the NP equation (**Equation 5**)^45^. An equivalent form of the NP equation can be found according to the divergence theorem (**Equation 5**)^46^, so that the number (*n*_*p*_) and area (*A*_*p*_) of electropores can be considered in the analysis. The flux term (*J*) incorporates the effects of concentration (*c*), fluid velocity (*v*), electric field (*E*), temperature (*T*), and cargo diffusivity in water at 37°C (*D*_*AB*_). The electric field is related to flux using the molecule charge (z), elemental charge (e), and Boltzmann constant (k_b_).

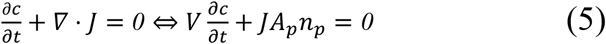

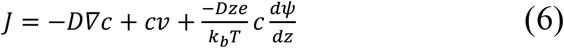

The flux of the molecules (*J*) is the vector sum of contributions from diffusion, advection, and electromigration (**Equation 6**). The equation was simplified by removing the effect of advection (there is no fluid velocity) and calcein diffusion. Molecular diffusion was neglected since the calculated electromotive species flux across the nanoporous membrane was ∼1350× greater than that of diffusion (**Supplemental Information S4)** and dominates cargo transport. The flux is then plugged into **Equation 5** with no assumed reaction and a transient concentration of cargo inside the cell volume (*V*). Here, *A*_*p*_ represents the area of one electropore on the cell and *n*_*p*_ is the number of electropores. When the simplified **Equation 6** is plugged into **Equation 5** and integrated with respect to time, it yields **Equation 7**.

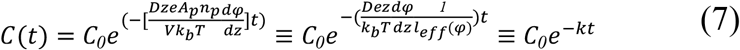

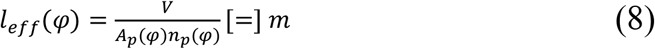

We further lump the electropore area, number of electropores, and the cell volume into a fitting parameter *l*_*eff*_, as these values can be difficult to ascertain or are uncertain during the experiment (**Equation 8**). As discussed in the main text, this yields an interpretable meaning for the experimental calcein rate constants. We use this parameter to adjust the timescale of our simulation equation to match experimental data by fitting *l*_*eff*_ to minimize the mean absolute error. Specific values used in the simulation are included in **Supplemental Information S4**.

This lumped model for charged molecule transport is mapped onto our device geometry by utilizing the electric field at each point in space retrieved from COMSOL simulations in a 3-D NanoEP device. Our NP simulation is 0-D and treats the hypothetical cell as a point in space, but we use input voltages from different (x, y) points in the COMSOL solution space to effectively predict calcein depletion rates over the entire membrane geometric area.

### Evaluation of Cargo Depletion and Decay in Rate Constant/Intensity Distance from Edge/Corner

Cargo depletion was assessed by tracking the reduction in fluorescent signal intensity of cells pre-stained with calcein AM during time-lapse imaging, conducted using the ECHO Revolve microscope. The fluorescent intensity drop is modeled by exponential decay in **Equation 1**. *I*_*f*_ is the fluorescence intensity of each pixel (or bin) tracked over time (t), starting at an initial fluorescence intensity *I*_*f,o*_ and reaching a plateau (*b*). Depletion rate constants (*k*) were extracted by fitting the experimental data to **Equation 1**^28^. The normalized depletion rate as a function of distance for the angled geometries (60°C, 90°C, and 120°C NanoEP devices) is also modeled by exponential decay in **Equation 1** for determining the characteristic length and Δ*k*_*norm*_ for the rate constant heat maps and the delivery of BSA A647 in the triangle, however rather than a change over time, the change would occur over a distance. Time-lapse imaging data were analyzed using Python 3.11.5 with the packages cv2, matplotlib, seaborn, scipy, and sklearn. Exponential regression models were optimized for each pixel by minimizing the sum of squared errors. The depletion rate constants were visualized as a heat map, where each pixel directly corresponds to the same location in the sample, and the color intensity reflects the value of the depletion rate constant. For the binned calcein depletion data, MATLAB was used to measure the fluorescence intensity decrease over time in each bin.

