## Supplemental Information for "Membrane Geometric Confinement Reshapes the Lateral Electric Field Distribution and Intracellular Cargo Transport in Nanopore Electroporation"

### Table of Contents


| Supplementary Information | Page Number |
|---|---|
| S1: Edge Effect Observed in Various Electroporation Buffers | 3 |
| S2: Calcein Signal Intensity vs Time During Electric Pulses | 3 |
| S3: Calcein Signal Intensity vs Time Without Electric Pulses | 4 |
| S4: Calculation of Flux Contributions For Nernst-Planck Equation | 5 |
| S5: Derivation of Cargo Transport Rate as a Function of Electropore Area | 6 |
| S6: EDAX Mapping of Oxygen Content in ITO After Electric Pulses | 7 |
| S7: Emergence of Edge Effect: Interplay of EEI and PCTE Geometry | 8 |
| S8: Charge Transfer Resistance ($R_{ct}$) Assumptions | 8 |
| S9: Angled Geometries: Images Before & After Calcein Removal | 9 |
| S10 Angled Geometries: Rate Comparison | 9 |
| S11: PDMS Pillars for Even Cargo Distribution During Delivery | 10 |
| S12: Delivery of BSA-A647 in 60°Angle Geometry | 10 |
| S13: Comparing Flux Pattern of Cargo Delivery & Depletion in 60° Angle | 11 |
| S14: Circle vs Serp: Comparison of Depletion Patterns | 12 |
| S15: Circle vs Serp: Comparison of Cargo Delivery Intensities | 13 |
| S16: Circle vs Serp: Comparison of Cargo Delivery Patterns | 14 |
| S17: Circle vs Serp: Comparison of BSA-AF647 Delivery Patterns | 15 |
| S18: Circle vs Serp: Comparison of Plasmid+YOYO-1 Delivery Patterns | 16 |
| S19: Purification of Free YOYO-1 Molecules from Bound Plasmid+YOYO-1 | 17 |

### S1: Edge Effect Observed in Various Electroporation Buffers

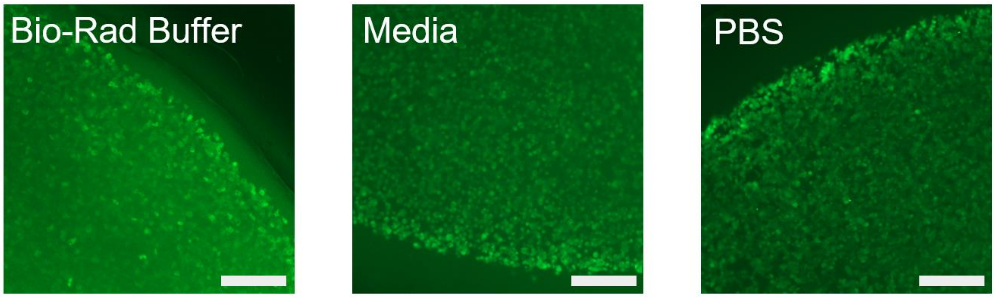


**Figure S1:** Edge effect observed in various electroporation buffers for BSA-AF488 delivery in HT1080 cells (20 V, 20 Hz, 1 ms square-wave pulse width, 10 s duration). Left: Gene Pulser Electroporation Buffer (Bio-Rad). Middle: Eagle's Minimal Essential Medium (EMEM) with 10% fetal bovine serum and 1% pen-strep. Right: Dulbecco's Phosphate Buffered Saline (DPBS) (scale bars, 300 $\mu$m).

### S2 & S3: Calcein Signal Intensity: Electric Pulses vs Photobleaching

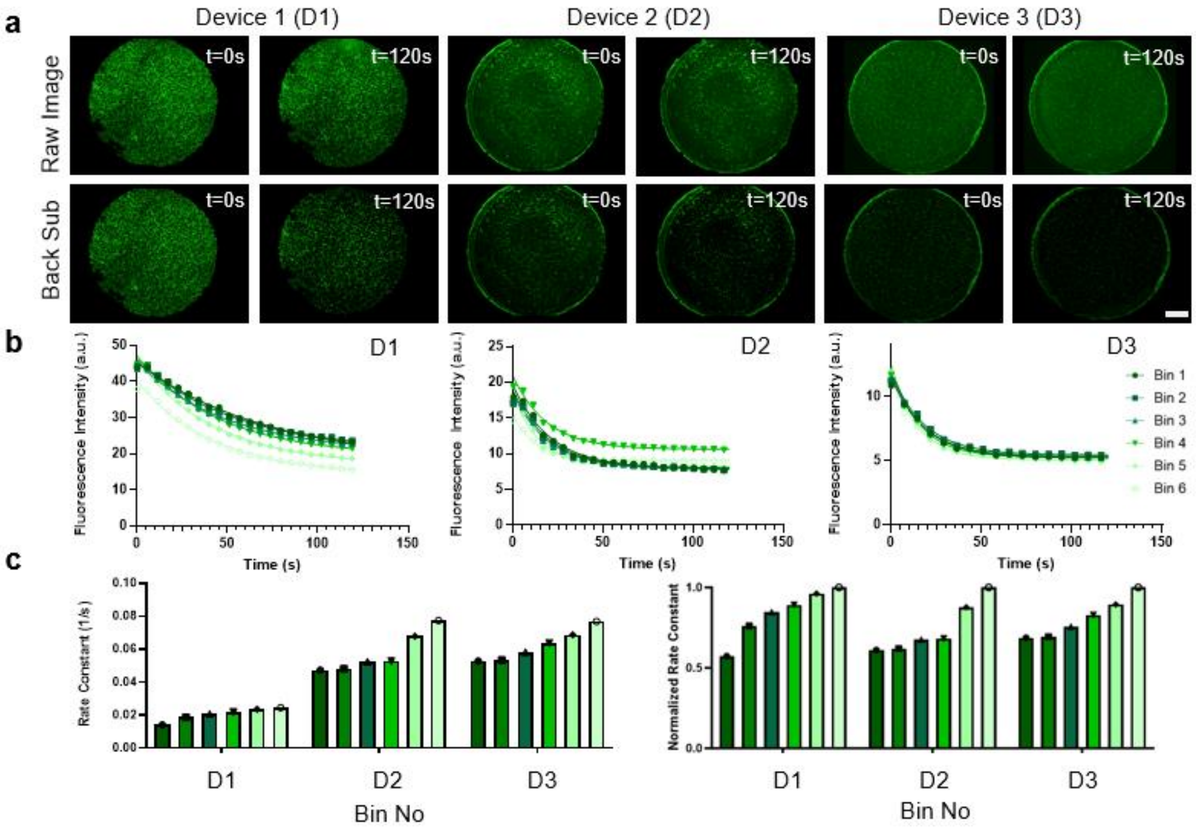


**Figure S2: a,** Before and after fluorescent micrographs of calcein depletion of HT1080 cells (15 V, 20 Hz, 1 ms square-wave pulse width, 120 s duration). Top: before and after fluorescence micrographs of raw images of full-device. Bottom: before and after fluorescence micrographs of background subtracted images of full-device (scale bar, 1 mm). **b,** Binned fluorescence intensity decrease of calcein over time with exponential fit (15 V, 20 Hz, 1 ms square-wave pulse width,

120 s duration; $n$ = 3; Bin 1 = center; Bin 6 = edge). Average $R^2$ values for exponential fits of radial bins were 0.991 +/- 0.004. **c,** Normalized rate constants from experimental data across binned regions. Each rate constant bin value was divided by the raw rate constant in Bin 6 for normalization.

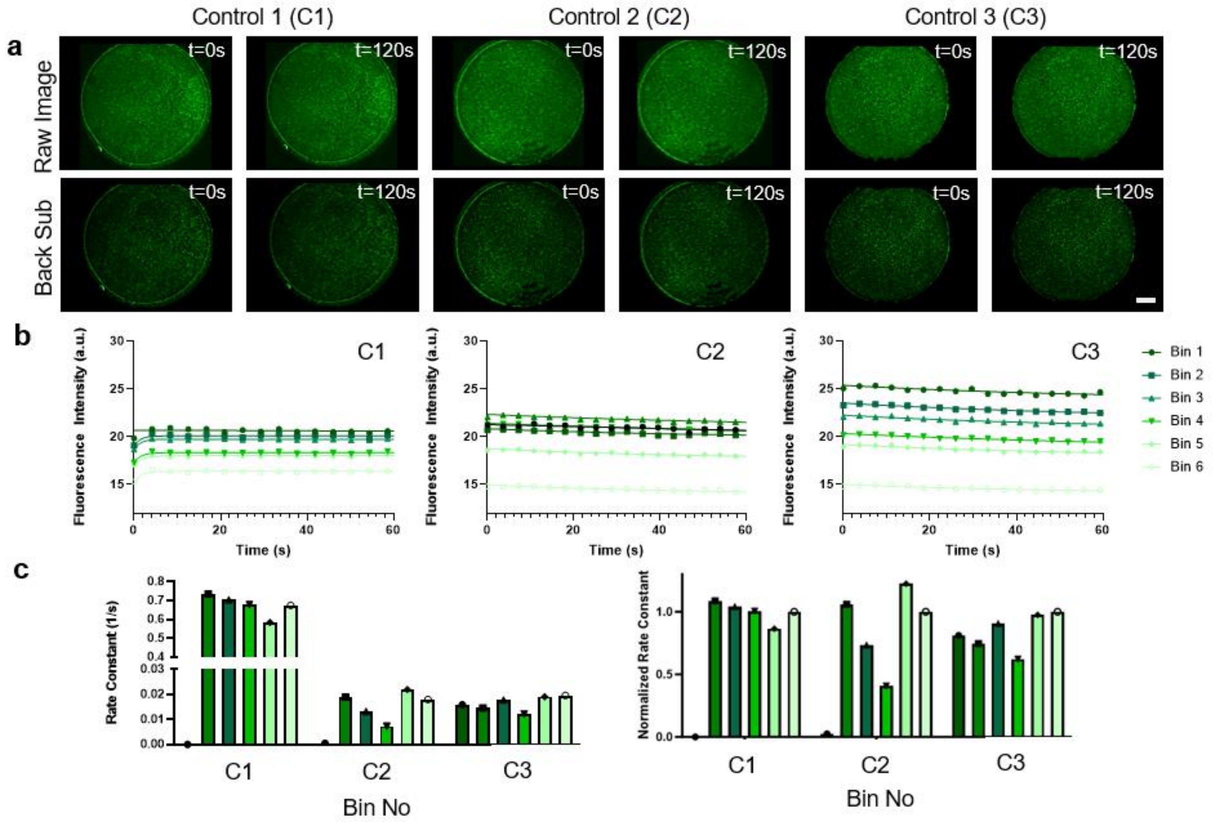


**Figure S3: a,** Before and after fluorescent micrographs of controls for calcein depletion experiments with no pulses applied. Top: before and after fluorescence micrographs of raw images of full-device. Bottom: before and after fluorescence micrographs of background subtracted images of full-device (scale bar, 1 mm). **b,** Binned fluorescence intensity as a function of time with exponential fit (Bin 1 = center; Bin 6 = edge). Average $R^2$ values for exponential fits of radial bins were 0.786 +/- 0.131. **c,** Normalized rate constants from experimental data across binned regions. Each rate constant bin value was divided by the raw rate constant in Bin 6 for normalization.

### S4. Calculation of Flux Contributions For Nernst-Planck Equation

Nernst-Planck Flux term: $J = -D\nabla c + \frac{Dze}{k_bT} c \frac{d\psi}{dz}$

Diffusion: $-D\nabla c$

- $D = 1 \cdot 10^{-10}$ m$^2$/s (calcein diffusion at 37 C) [cite]
- $\nabla c = \frac{dc}{dz} = \frac{0.005 \frac{mol}{m^3} - 0 \frac{mol}{m^3}}{12 \cdot 10^{-6} m} = 416.67 \frac{mol}{m^4}$
- $D\nabla c = 4.16 \cdot 10^{-8} \frac{mol}{m^2 s}$

Electromigration: $\frac{Dze}{k_bT} c \frac{d\psi}{dz}$

- $D = 1 \cdot 10^{-10}$ m$^2$/s (calcein diffusion at 37 C)
- $z = 3$ (assumed calcein charge at pH 7.2)
- $e = 1.602 \cdot 10^{-19}$C
- $k_bT = 4.28 \cdot 10^{-21}$J
- $c = 0.005 \frac{mol}{m^3}$
- $\frac{d\psi}{dz} = \frac{12\,V}{12 \cdot 10^{-6} m} = 1 \cdot 10^6 \frac{V}{m}$
- $\frac{-Dze}{k_bT} c \frac{d\psi}{dz} = 7.49 \cdot 10^{-5} \frac{mol}{m^2 s}$

Ratio: $\frac{Dze}{k_bT} c \frac{d\psi}{dz} \,/\, D\nabla c = 1347.48$

**Supplemental Information S4**: To leverage simulated voltage profiles to easily predict cargo depletion rates, a simplified Nernst–Planck (NP) equation was implemented in Python. The movement of charged molecules in solution is described by the NP equation. There is no bulk fluid flow in our system, and so the advection term is naturally neglected. To determine if we could neglect the diffusion term as well, we computed the ratio of the electromigration term and the diffusion term, and we found the electromigration term to be >3 orders of magnitude larger. Thus, we could avoid partial differential treatments and simplify our empirical model.

### S5: Derivation of Cargo Transport Rate as a Function of Electropore Area

1. $\frac{\partial c}{\partial t} + \nabla \cdot J = 0$
2. $J = \frac{-Dze}{k_b T} c \frac{d\psi}{dz}$
3. $\frac{\partial c}{\partial t} + \frac{Dze}{k_b T}\left(\frac{\partial c}{\partial z}\frac{d\psi}{dz} + c\frac{d^2\psi}{d^2 z}\right) = 0 \Rightarrow$

$$\frac{\partial c}{\partial t} = \frac{-Dze}{k_b T}\frac{\partial c}{\partial z}\frac{d\psi}{dz}; \frac{d^2\psi}{d^2 z} = 0; \frac{\partial c}{\partial z} = \frac{c}{l}$$

4. $\frac{1}{c}\partial c = \frac{-Dze}{k_b T}\frac{1}{l}\frac{d\psi}{dz}\partial t$
5. $ln(c) = \frac{-Dze}{k_b T}\frac{1}{l}\frac{d\psi}{dz} t + C \Rightarrow$

$$c = c_o e^{\frac{-Dze}{k_b T}\frac{1}{l}\frac{d\psi}{dz} t} = c_o e^{-k_c t}$$

1. $V\frac{\partial c}{\partial t} + J A_p n_p = 0$
2. $J = \frac{-Dze}{k_b T} c \frac{d\psi}{dz}$
3. $V\frac{\partial c}{\partial t} + \frac{Dze}{k_b T} c \frac{d\psi}{dz} A_p n_p = 0 \Rightarrow \frac{\partial c}{\partial t} = \frac{-Dze}{k_b T} c \frac{d\psi}{dz}\frac{A_p n_p}{V}$
4. $\frac{1}{c}\partial c = \frac{-Dze}{k_b T}\frac{A_p n_p}{V}\frac{d\psi}{dz}\partial t$
5. $ln(c) = \frac{-Dze}{k_b T}\frac{A_p n_p}{V}\frac{d\psi}{dz} + C \Rightarrow$

$$c = c_o e^{\frac{-Dze}{k_b T}\frac{A_p n_p}{V}\frac{d\psi}{dz} t} = c_o e^{-k_c t}$$

**Supplemental Information S5**: Derivation of the intracellular cargo concentration under cargo depletion conditions. The left and right side are equivalent formulations of the Nernst-Planck continuity equation. The final forms of the equations are similarly equivalent, and with an exponentially decaying form. Upon inspection, they have identical terms in the exponent except for $1/l$ and $A_p n_p / V$. These two terms have equivalent units (1/m), and we utilized this derivation to create the empirical parameter of *effective length scale* ($l_{eff}$).

### S6: EDAX Mapping of Oxygen Content in ITO After Electric Pulses

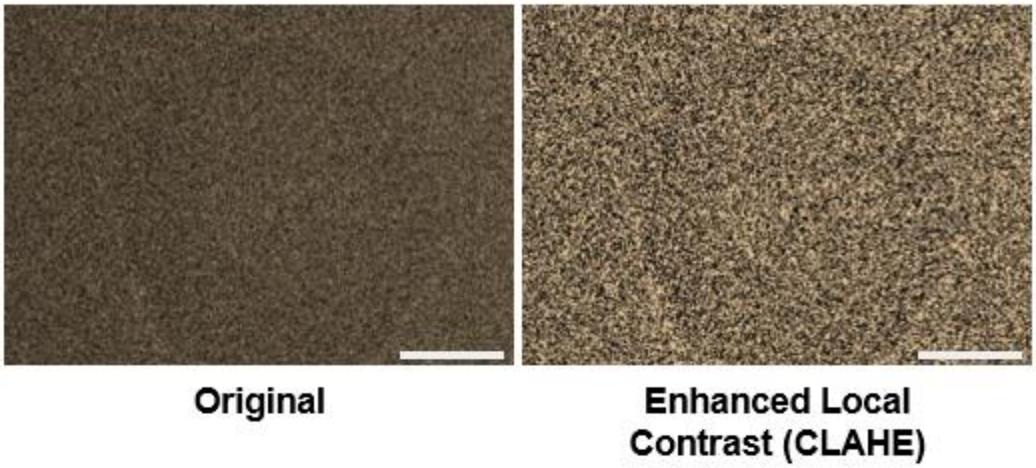


**Figure S6:** Energy Dispersive X-Ray elemental map of oxygen on a portion of ITO glass that underwent electrochemical reduction. Brighter colors indicate more oxygen. (scale bars, 500 $\mu$m). SEM imaging was performed using the Thermo Fisher Nova 200 Nanolab at the Michigan Center for Materials Characterization ($MC^2$). The ITO coated glass was mounted using conductive carbon tape and grounded using colloidal graphite glue (Electron Microscopy Sciences). Images of ITO surfaces were acquired using secondary electrons at 5 kV and 1.6 nA of beam current. EDX maps were collected at 5 kV and 3.2 nA of beam current, such that the electron plume would not probe too deep beneath the surface.

### S7 & S8: Emergence of Edge Effect: Interplay of EEI and PCTE Geometry

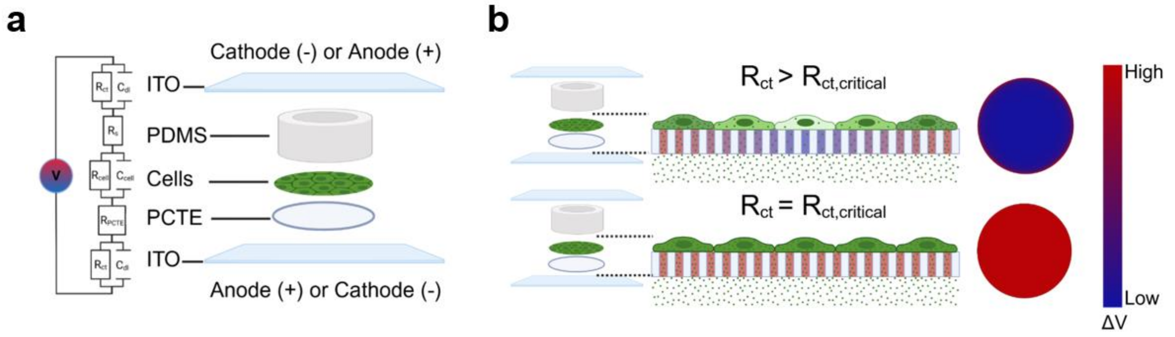


**Figure S7: a)** Left: simplified equivalent circuit diagram of NanoEP system ($R_{ct}$ is the charge transfer resistance, $C_{dl}$ is the double layer capacitance, $R_{PCTE}$ is the resistance of the PCTE membrane, $R_{cell}$ and $C_{cell}$ are the resistance and capacitance of the cell monolayer respectively, and $R_s$ is the solution resistance). Right: Schematic illustration of the NanoEP device used for cargo delivery and depletion experiments. **b)** Schematic illustration of the voltage drop across the PCTE membrane at two different $R_{ct}$ values. $R_{critical}$ is defined as the $R_{ct}$ value where the center-to-edge variation in electrolyte voltage drop is <500 mV. Top: edge effect is observed when $R_{ct}>R_{ct,critical}$. This includes both a higher voltage drop and an increase in cargo delivery near the edge of the device as compared to the center (side view). COMSOL simulation showing higher voltage drop at periphery of device compared to center (top view). Bottom: edge effect is not observed when $R_{ct}=R_{ct,critical}$. Even cargo delivery and voltage drop at edge and center of device (side view). COMSOL simulation showing even voltage drop at periphery and center of the device (top view).

**Supplemental Information S8: Charge Transfer Resistance ($R_{ct}$) Assumptions**

In our simulations, charge transfer resistance ($R_{ct}$) was made constant across the EEI interface; however, $R_{ct}$ is an exponentially decaying function of voltage. This means that the larger driving force at the edge should lower $R_{ct}$ locally, further enhancing the z-directional voltage drop across the nanopore membrane. Practically though, our system is operating at much higher voltages than those relevant to electrochemical studies (>10 V). This high voltage causes large overpotentials for the assumed reactions (ITO reduction, hydrogen evolution, oxygen evolution[32,47], thus Rct is expected to be very low ($\ll 1\ \Omega$)[47] and unlikely to affect the trends we show here. Through this analysis, we validated a difference in the vertical electric field around the device through material property changes and how it extends to NanoEP cargo flux.

### S9 & S10: Angled Geometry & the Effect on Calcein Depletion

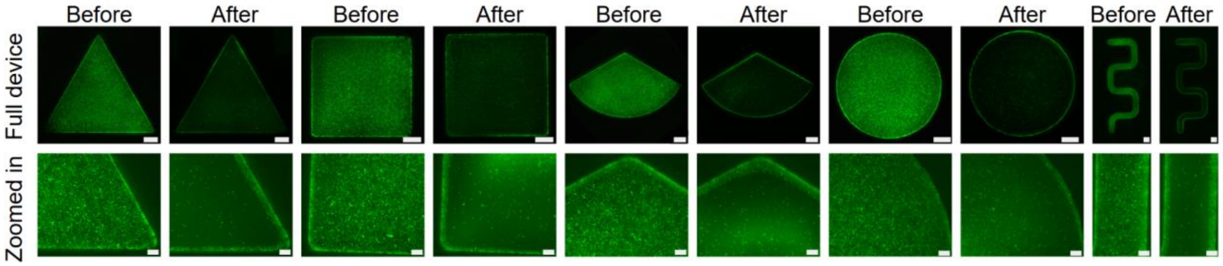


**Figure S9:** Before and after representative fluorescent micrographs of calcein depletion of HT1080 cells (15 V, 20 Hz, 1 ms square-wave pulse width, 120 s duration; $n = 3$). Top: full-device before and after fluorescence micrographs (scale bars, 1 mm). Bottom: Zoomed in before and after fluorescence micrographs (scale bars, 200 $\mu$m). We attached the PCTE to the PDMS device using uncured PDMS as an adhesive, and wet PDMS will wick over the adjacent, uncovered PCTE membrane area. Due to this, the cells adhered directly next to the PDMS wall do not lose fluorescence in basically every case.

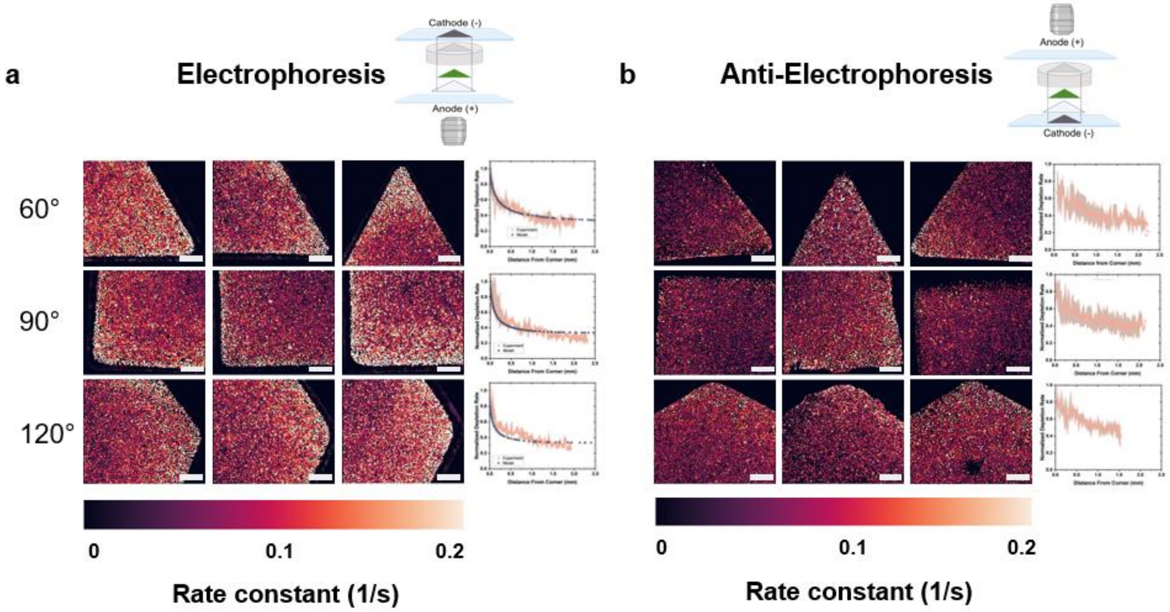


**Figure S10: a)** Left: experimental rate constant heat maps of electrophoretic calcein depletion for 60°, 90°, 120° NanoEP devices (scale bars, 430 $\mu$m). Right: Comparison of simulated and experimental electrophoretic normalized rate constants as a function of distance from corner for 60°, 90°, and 120° NanoEP devices ($n = 3$). **b)** Left: experimental normalized rate constant heat maps of anti-electrophoretic calcein depletion for 60°, 90°, 120° NanoEP devices (scale bars, 430 $\mu$m). Right: experimental anti-electrophoretic normalized rate constants as a function of distance from corner for 60°, 90°, and 120° NanoEP devices ($n = 3$).

### S11: PDMS Pillars for Even Cargo Distribution During Delivery

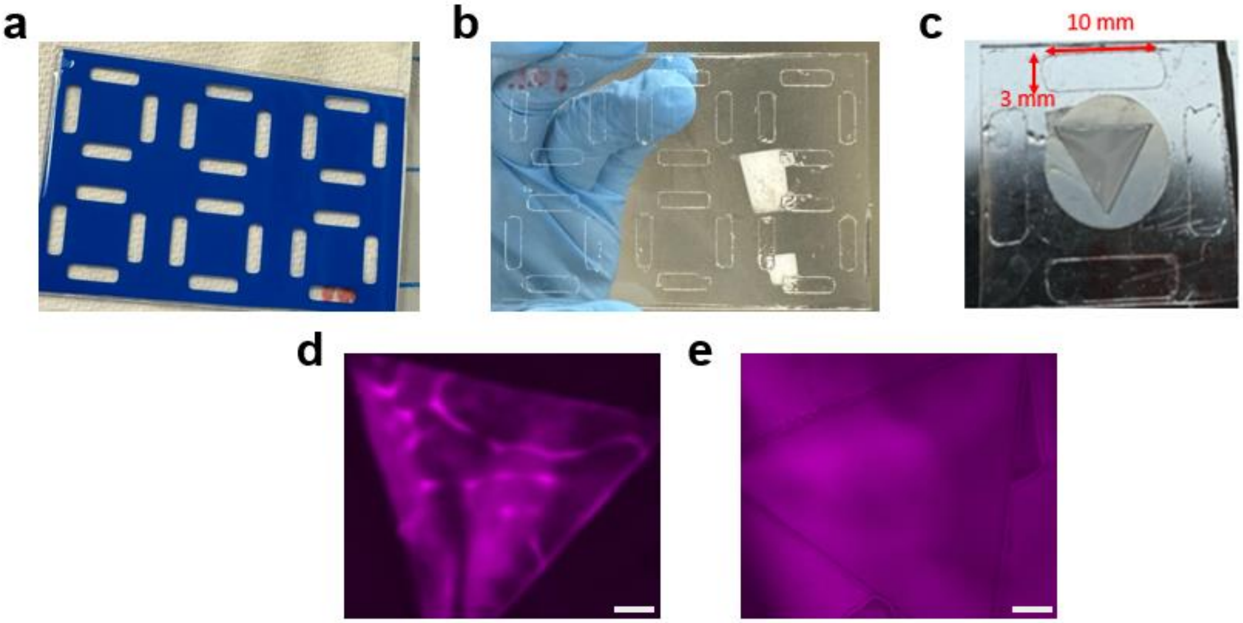


**Figure S11: a)** Sticker with pillars (made using Scancut Brother Cutting Machine) attached to a glass slide. PDMS was spun coat on a glass slide to create a 200 $\mu$m layer. The glass slide was then cured overnight at 60°C. **b)** The sticker was removed so only the pillars remained. **c)** Pillars were then plasma bonded or attached to the NanoEP devices with uncured PDMS (followed by an overnight cure at 60°C). **d)** BSA-AF647 localization from folds of the PCTE membrane with no pillars underneath the membrane (scale bar, 1 mm). **e)** BSA-AF647 spreading with 200 $\mu$m tall pillars underneath membrane (scale bar,  1 mm).

### S12: Delivery of BSA-A647 in 60°Angle Geometry

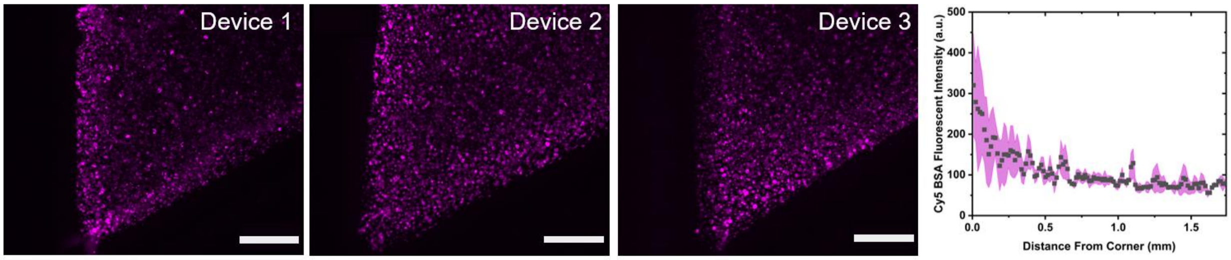


**Figure S12: a)** Fluorescence micrographs of 60° corner of BSA-AF647 delivery (20 V, 20 Hz, 1 ms square-wave pulse width, 10 s duration  ($n$ = 3; scale bar, 430 $\mu$m). Fluorescence intensity profile of BSA-AF647 delivery as a function of distance from a 60° device corner.

### S13: Comparing the Flux Patterns of Cargo Delivery & Depletion: 60° Angle Geometry

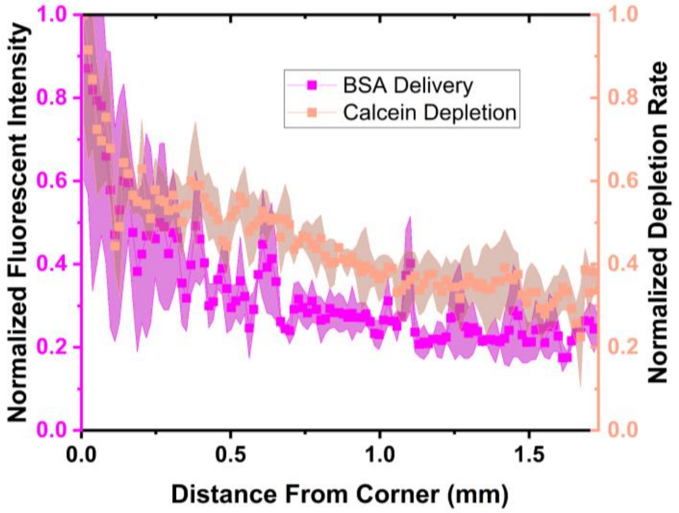


**Figure S13:** Comparison of the normalized calcein depletion rate constants and the normalized fluorescent intensity of BSA-AF467 delivery from **Figure 3** as a function of distance from the corner for 60° NanoEP devices (*n=3*).

### S14: Circle vs Serpentine: Comparison of Depletion Patterns

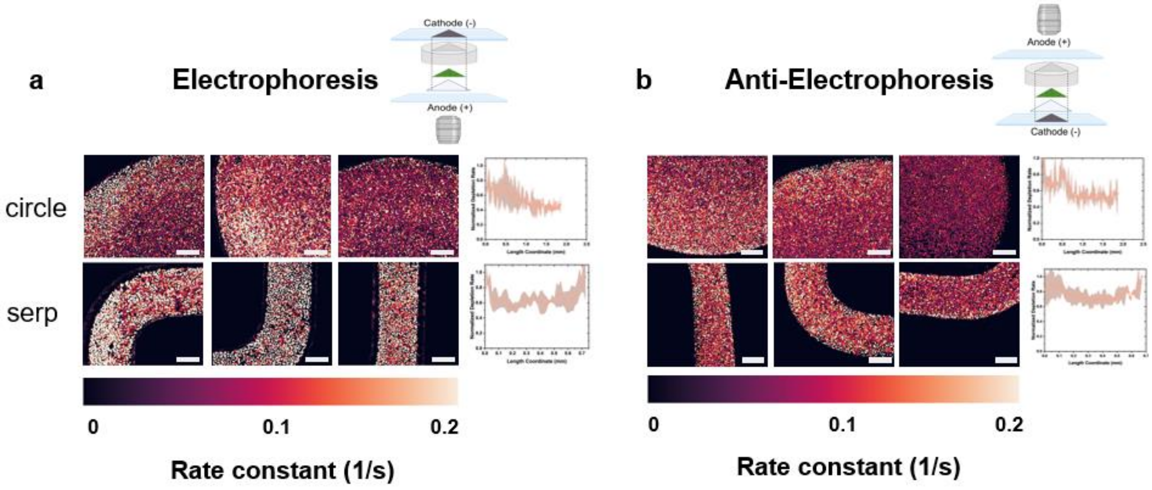


**Figure S14: a)** Left: experimental rate constant heat maps of electrophoretic calcein depletion for serpentine and circle NanoEP devices (scale bars, 430 $\mu$m). Right: Comparison of experimental electrophoretic normalized rate constants as a function of distance from corner for serpentine and circle NanoEP devices ($n$ = 3). **b)** Left: experimental rate constant heat maps of anti-electrophoretic calcein depletion for serpentine and circle NanoEP devices (scale bars, 430 $\mu$m). Right: Comparison of experimental anti-electrophoretic normalized rate constants as a function of distance from corner for serpentine and circle NanoEP devices ($n$ = 3).

### S15: BSA-A647 Delivery, Plasmid+YOYO-1 Delivery, and DsRed Expression in Circle and Serpentine

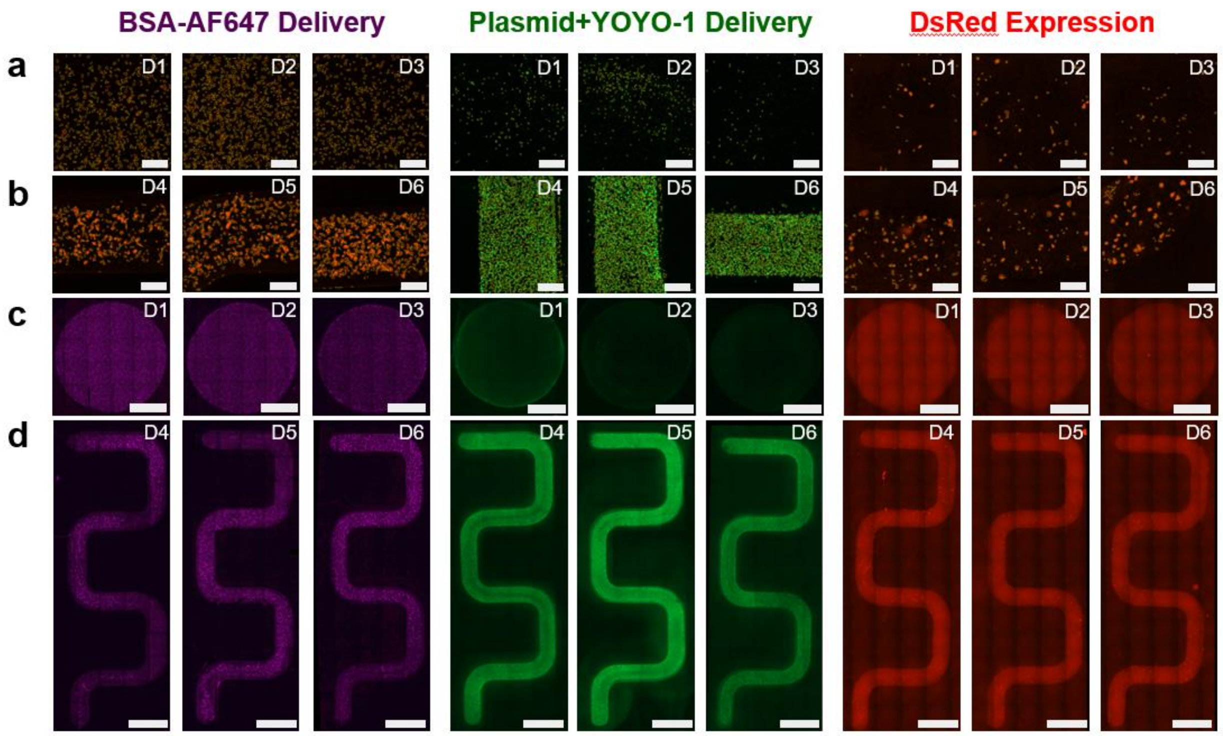


**Figure S15:** Left: Delivery of BSA-AF647 (2.5 mg/mL; 20 V, 20 Hz, 1 ms square-wave pulse width, 10 s duration) in HT1080 cells in circle and serpentine geometry. Middle: Delivery of Plasmid+YOYO-1 (100 ng/$\mu$L pLenti3.7-DsRed plasmid; 25 V, 1Hz, 10 ms square-wave pulse width, 4 s duration) in HT1080 cells in circle and serpentine geometry. Right: Expression of DsRed (5 ng/$\mu$L pLenti3.7-DsRed plasmid; 25 V, 1 Hz, 10 ms square-wave pulse width, 4 s duration). $n = 3$ for each geometry in each experiment (BSA-AF647, Plasmid+YOYO-1, and DsRed expression). **a)** Representative fluorescent micrographs for each circle device geometry with cell recognition software identifying cells with yellow boundary (scale bars, 300 $\mu$m). **b)** Representative fluorescent micrographs for each serpentine device geometry with cell recognition software identifying cells with yellow boundary (scale bars, 300 $\mu$m). **c)** Fluorescent micrographs of full circular devices (scale bars, 2 mm). **d)** Fluorescent micrographs of full serpentine devices (scale bars, 2 mm).

### S16: Circle vs Serpentine: Comparison of Cargo Delivery Patterns

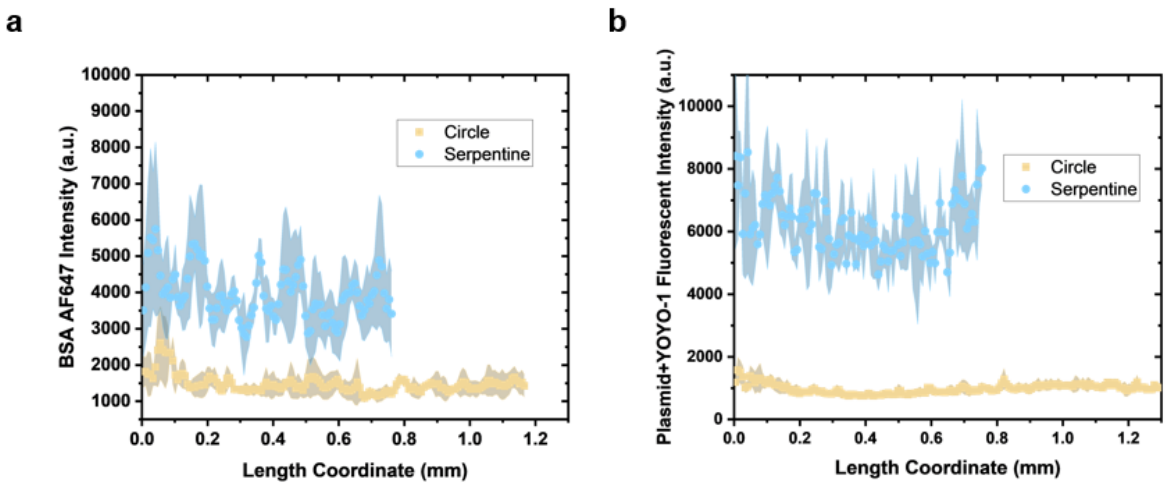


**Figure S16: a)** Comparison of BSA-AF647 intensity as a function of distance from the edge for serpentine vs. circular devices ($n$ = 3 per geometry). **b)** Comparison of plasmid labeled with YOYO-1 intensity as a function of distance from the edge for serpentine vs. circular devices ($n$ = 3 per geometry).

### S17: Circle vs Serpentine: Comparison of BSA-AF647 Delivery Patterns

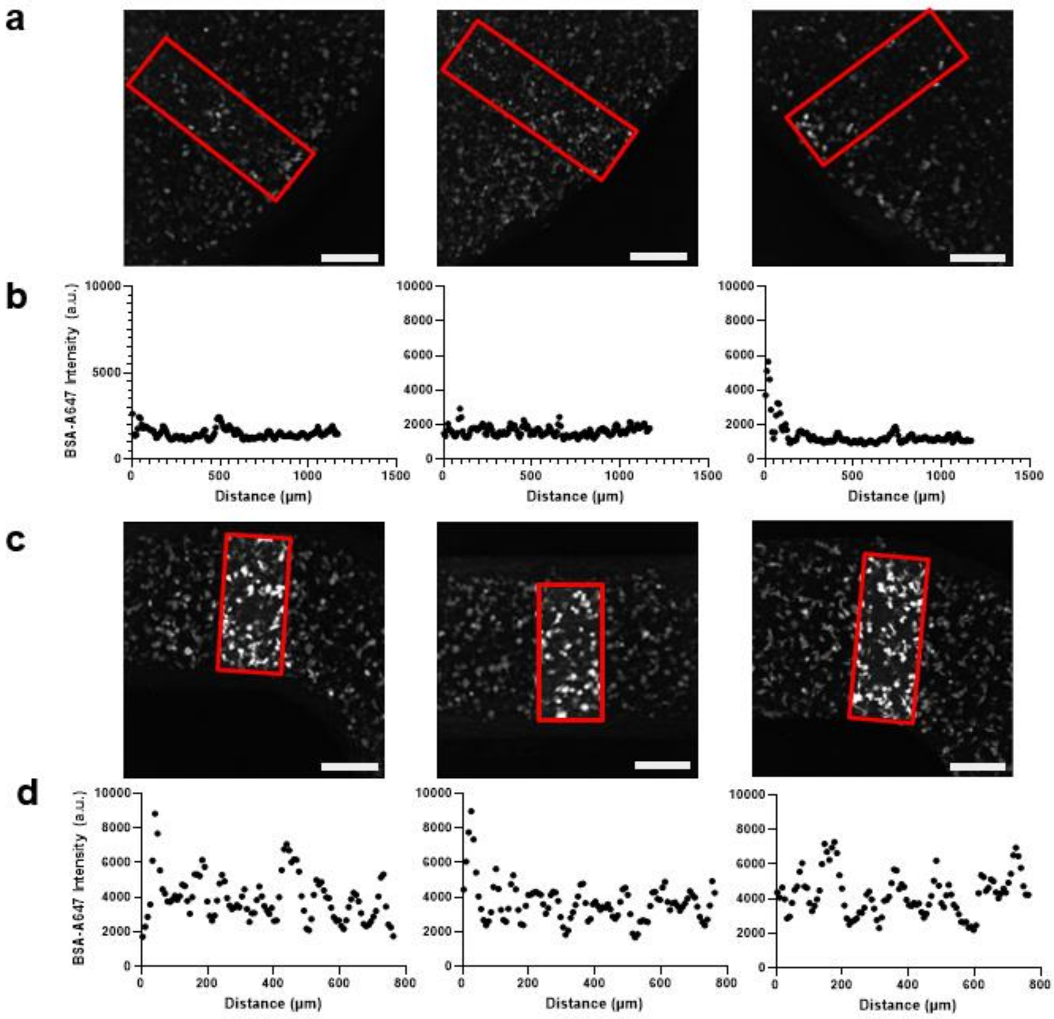


**Figure S17: a)** Fluorescent micrographs of BSA-A647 delivery in circle geometries with region of interest for line plot analysis (370 $\mu$m thick line drawn from edge towards center; scale bars, 300 $\mu$m). **b)** BSA-AF647 intensity profiles as a function of distance from the edge. Each intensity value on the plot corresponds to the average intensity in a 7.5 $\mu$m binned region along region of interest. **c)** Fluorescent micrographs of BSA-A647 delivery in serpentine geometries with region of interest for line plot analysis (370 $\mu$m thick line drawn from edge towards center; scale bars, 300 $\mu$m). **d)** BSA-AF647 intensity profiles from edge to edge in serpentine device. Each intensity value on the plot corresponds to the average intensity in a 7.5 $\mu$m binned region along region of interest.

### S18: Circle vs Serpentine: Comparison of Plasmid+YOYO-1 Delivery Patterns

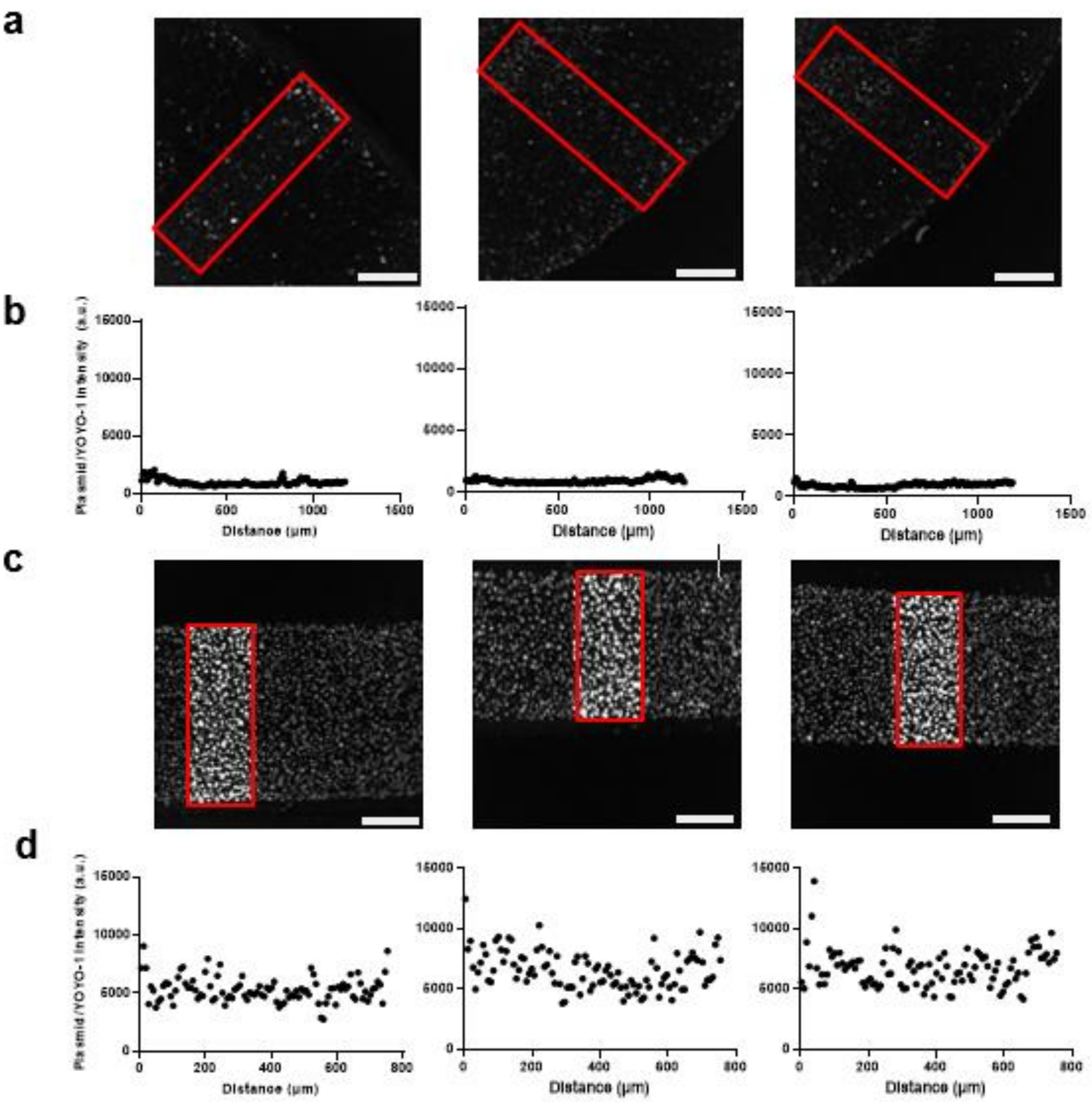


**Figure S18: a)** Fluorescent micrographs of Plasmid+YOYO-1 delivery in circle geometries with region of interest for line plot analysis (370 $\mu$m thick line drawn from edge towards center; scale bars, 300 $\mu$m). **b)** Plasmid+YOYO-1intensity profiles as a function of distance from the edge. Each intensity value on the plot corresponds to the average intensity in a 7.5 $\mu$m binned region along region of interest. **c)** Fluorescent micrographs of Plasmid+YOYO-1 delivery in serpentine geometries with region of interest for line plot analysis (370 $\mu$m thick line drawn from edge towards center; scale bars, 300 $\mu$m). **d)** Plasmid+YOYO-1 intensity profiles from edge to edge in serpentine device. Each intensity value on the plot corresponds to the average intensity in a 7.5 $\mu$m binned region along region of interest.

### S19: Purification of Free YOYO-1 Molecules from Bound Plasmid+YOYO-1

Two samples, containing plasmid+YOYO-1 or YOYO-1 only (both diluted in elution buffer and containing the same amount of YOYO-1 molecules), were run through a PCR purification kit. The purpose was to remove any free YOYO-1 molecules that were not bound to the plasmid. After purification, the two samples were delivered into the NanoEP devices at 20 V, 1 Hz, 10 ms square-wave pulses for 8 s duration. There was no positive YOYO-1 signal observed in the purified YOYO-1 only sample. We observed positive YOYO-1 signal in the purified plasmid+YOYO-1 sample. Thus, for plasmid+YOYO-1 delivery experiments, we assumed after filtering the plasmid+YOYO-1 in the purification kit, the positive YOYO-1 signal in the cells post electroporation also correlated to plasmid delivery into the cell. Since we diluted the plasmid+YOYO-1 in the electroporation buffer, we also analyzed the intensity change using the same dilution in the electroporation buffer compared to the elution buffer. We observed a 22-29% reduction in fluorescence intensity in the electroporation buffer compared to the elution buffer. However, most of the fluorescence is still retained in the electroporation buffer.

**a**

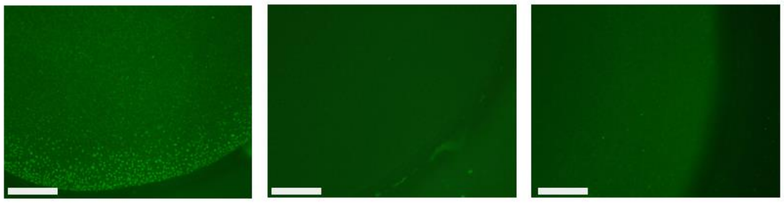

| | Fold Increase Relative to Elution Buffer |
|---|---|
| Purified Plasmid/YOYO-1 | 145 |
| Purified YOYO-1 | 1 |

**b**

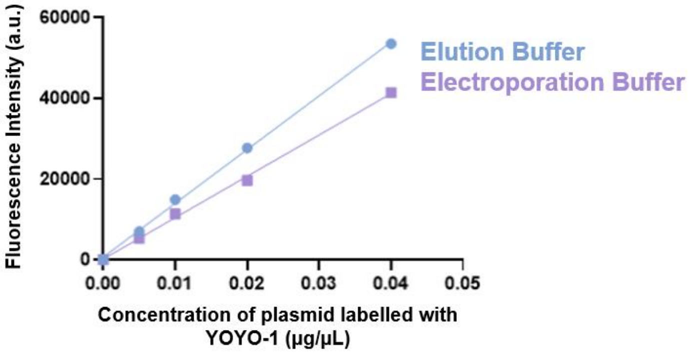


**Figure S19: a)** Left to middle right: (left) delivery of purified plasmid labelled with YOYO-1 (20 V, 1 Hz, 10 ms square-wave pulse width, 8s duration), (middle left) purified YOYO-1 with no plasmid (20 V, 1 Hz, 10 ms square-wave pulse width, 8s duration), (middle right) purified plasmid labelled with YOYO-1 with no pulses (scale bars, 430 $\mu$m), and (right) fold changes of purified plasmid labelled with YOYO-1 and purified YOYO-1 as compared to elution buffer. **b)** Comparison of fluorescence intensity between plasmid labeled with YOYO-1 in elution buffer and electroporation buffer.
